# *Borrelia burgdorferi* engages mammalian type I interferon responses via the cGAS-STING pathway

**DOI:** 10.1101/2022.05.13.491896

**Authors:** Lauren C. Farris, Sylvia Torres-Odio, L. Garry Adams, A. Phillip West, Jenny A. Hyde

## Abstract

*Borrelia burgdorferi*, the etiologic agent of Lyme disease, is a spirochete that modulates numerous host pathways to cause a chronic, multi-system inflammatory disease in humans. *B. burgdorferi* infection can lead to Lyme carditis, neurologic complications, and arthritis, due to the ability of specific borrelial strains to disseminate, invade, and drive inflammation. *B. burgdorferi* elicits type I interferon (IFN-I) responses in mammalian cells and tissues that are associated with the development of severe arthritis or other Lyme-related complications. However, the innate immune sensors and signaling pathways controlling IFN-I induction remain unclear. In this study, we examined whether intracellular nucleic acid sensing is required for the induction of IFN-I to *B. burgdorferi*. Using fluorescence microscopy, we show that *B. burgdorferi* associates with mouse and human cells in culture and we document that internalized spirochetes co-localize with the pattern recognition receptor cyclic GMP-AMP synthase (cGAS). Moreover, we report that IFN-I responses in mouse macrophages and murine embryonic fibroblasts are significantly attenuated in the absence cGAS or its adaptor Stimulator of Interferon Genes (STING), which function to sense and respond to intracellular DNA. Longitudinal in vivo tracking of bioluminescent *B. burgdorferi* revealed similar dissemination kinetics and borrelial load in C57BL/6J wild-type, cGAS-deficient, or STING-deficient mice. However, infection-associated tibiotarsal joint pathology and inflammation were modestly reduced in cGAS-deficient compared to wild-type mice. Collectively, these results indicate that the cGAS-STING pathway is a critical mediator of mammalian IFN-I signaling and innate immune responses to *B. burgdorferi*.

**KEY POINTS:**

- *B. burgdorferi* triggers type I interferon responses in macrophages and fibroblasts
- Coiled spirochetes are observed in the cytosol and co-localize with cGAS
- cGAS and STING mediate *B. burgdorferi*-induced type I interferon responses

## INTRODUCTION

Lyme disease results from infection with the spirochetal bacterium *Borrelia burgdorferi* and presents as a multi-systemic inflammatory disease causing debilitating morbidity due to fatigue, malaise, severe arthritis, and cardiac and neurologic complications (1–4). It is the most common tick-borne disease in the United States and is a significant public health concern, with the CDC reporting 39,000 cases per year but insurance reports indicating prevalence greater than 470,000 cases per year (5–7). Antibiotic treatment is highly effective when administered shortly after the tick bite, yet antibiotic efficacy declines as the borrelial infection progresses (8, 9). Lyme disease occurs in stages of localized, disseminated, and chronic infection as the extracellular pathogen spreads from the site of the tick bite to secondary tissues, including the joints, heart, and central nervous system, causing Lyme arthritis, carditis, and neuroborreliosis respectively (1–3, 8). The development of arthritis, a characteristic symptom of late Lyme disease in North America, is associated with a robust innate immune response that includes induction of type I interferon (IFN-I) cytokines (10–12).

*B. burgdorferi* elicits a robust innate immune response, resulting in the secretion of pro-inflammatory cytokines and chemokines (13–20). In addition, *B. burgdorferi* triggers IFN-I responses in a wide array of human cells, mouse cells, and infected tissues (11–13, 19, 21–28). Evidence suggests that innate IFN-I signaling plays key roles in several aspects of *B. burgdorferi* pathology (29–33). Notably, IFN-I and resulting interferon-stimulated gene (ISG) signatures are linked to the development of more severe arthritis in experimental models, and also correlate with lingering neurocognitive symptoms of Lyme disease (12, 30, 32, 34). In addition, a recent study from Lochhead and colleagues observed that a robust interferon gene signature correlates with decreased expression of tissue repair genes in synovial lesion biopsies from patients with postinfectious, *B. burgdorferi*-induced Lyme arthritis (35). It is well-appreciated that innate immune sensing of the abundant *B. burgdorferi* lipoproteins via Toll-like receptor 2 (TLR2) triggers production of pro-inflammatory cytokines and chemokines in vitro and in vivo (14, 36–38). However, lipoprotein binding to TLR2 is not a robust inducer of IFN-I responses in most mouse and human cell types (39, 40). Studies using human cells have implicated nucleic acid sensing TLRs (TLR 7, 8, and 9) as regulators of IFN-I induction in human immune cells challenged with *B. burgdorferi* in vitro (13, 18, 19). In contrast, other reports have shown that *B. burgdorferi* can engage IFN-I responses in non-phagocytic fibroblasts and endothelial cells, which do not express a full complement of TLRs (25, 41). Moreover, *B. burgdorferi*-related ISG induction in murine macrophages is independent of the two primary TLR adaptor proteins MyD88 and TRIF (22, 23). Thus, TLR-mediated sensing of *B. burgdorferi* pathogen-associated molecular patterns does not appear to be the predominant trigger of IFN-I responses in mammalian cells during infection.

Innate immune pathways that sense cytosolic nucleic acids, such as the RIG-I-like receptor (RLR)-Mitochondrial Antiviral Signaling (MAVS) or Cyclic GMP-AMP synthase (cGAS)-Stimulator of Interferon Genes (STING) pathway, have emerged as key regulators of IFN-I production in both immune and non-immune cells (42–49). Cyclic GMP-AMP synthase (cGAS) is an intracellular DNA sensor that localizes to the mammalian cell cytoplasm and nucleus. Upon binding intracellular pathogen DNA, micronuclei, or mitochondrial DNA, cGAS generates the non-canonical cyclic dinucleotide, 2’3’-cyclic guanosine monophosphate-adenosine monophosphate (cGAMP), which binds STING. This results in the recruitment and activation of Tank-binding Kinase 1 (TBK1), leading to the phosphorylation of Interferon Regulatory Factor 3 (IRF3) for the induction of type I interferons (IFNα, β) and ISGs (42–44).

Although initially identified as an antiviral host defense pathway, the cGAS-STING pathway is also critical for induction of robust IFN-I responses to intracellular bacteria such as *Mycobacterium tuberculosis* and *Listeria monocytogenes* (45, 50, 51). Moreover, recent work has shown that the cGAS-STING pathway is essential for IFN-I production in response to multiple extracellular pathogens, including *Pseudomonas aeruginosa*, *Klebsiella pneumoniae*, and *Staphylococcus aureus* (52, 53). *B. burgdorferi,* predominantly an extracellular pathogen, is readily taken up by phagocytic cells and associates with endothelial cells and fibroblasts, which are key sources of IFN-I in the Lyme disease joint (54–62). *B. burgdorferi* also produces cyclic dinucleotides c-di-GMP and c-di-AMP, which can directly engage STING (63–66). Thus, there are several potential routes by which *B. burgdorferi* infection could trigger the cGAS-STING-IFN-I pathway.

In this study, we tested the hypothesis that *B. burgdorferi* infection induces IFN-I through the cGAS-STING pathway. We exposed phagocytic and non-phagocytic cells lacking various components of the cytosolic nucleic acid sensing machinery to viable and sonicated *B. burgdorferi* and evaluated IFN-I and ISG expression after exposure. We also assessed the degree of association of *B. burgdorferi* with cultured fibroblasts and examined co-localization of cGAS with intracellular spirochetes. Furthermore, we performed an infectivity study with bioluminescent *B. burgdorferi* in mice deficient in cGAS or STING to assess borrelial load by in vivo imaging and joint inflammation using histopathology. Our results reveal that *B. burgdorferi* engages IFN-I responses in a cGAS-STING dependent manner without significantly altering infection kinetics or borrelial load in tissues.

## MATERIALS AND METHODS

### Mouse and *B. burgdorferi* strains

C57BL/6J (strain 000664), cGAS deficient (cGAS^KO^, strain 026554), STING deficient (STING^KO^, strain 017537), IFNAR deficient (IFNAR^KO^, strain 028288) and MAVS deficient (MAVS^KO^, strain 008634) were obtained from the Jackson Laboratory. MAVS^KO^ mice were backcrossed to C57BL/6J mice for 10 generations before generation of primary cell lines. Mice were group-housed in humidity-controlled environments maintained at 22°C on 12-hour light– dark cycles (600–1800). Food and water were available ad libitum. All animal experiments were conducted in accordance with guidelines established by Department of Health and Human Services Guide for the Care and Use of Laboratory Animals and the Texas A&M University Institutional Animal Care and Use Committee.

Low-passage *B. burgdorferi* strains B31-A3 and ML23 pBBE22*luc* were cultured in BSK-II medium with 6% normal rabbit serum (NRS) (Pel-Freeze Biologicals, Rogers, AR) and grown to mid-log phase at 37°C at 5% CO_2_(67–71). ML23 pBBE22*luc* cultures were supplemented with 300 μg/ml kanamycin.

### Cell culture and *B. burgdorferi* infection

Primary mouse embryonic fibroblasts (MEFs) were generated from WT, cGAS^KO^, STING^KO^, IFNAR^KO^, and MAVS^KO^ E12.5 to E14.5 embryos. Cells were grown in DMEM (D5756, Millipore Sigma) containing 10% low endotoxin FBS (97068-085, VWR) and cultured for no more than four passages prior to experiments. SV40 immortalized cGAS^KO^ MEFs reconstituted with HA-tagged mouse cGAS were previously reported(72). Primary bone marrow-derived macrophages (BMDMs) were generated as described (73). Briefly, bone marrow cells were collected from the femur and tibia of mice and differentiated into macrophages in DMEM containing 10% low endotoxin FBS, and 20% (v/v) conditioned media harvested from L929 cells (CCL-1, ATCC). Cells were plated in Petri plates and maintained in L929-conditioned media for 7 days. The day before experiments, macrophages were plated in tissue culture plates and maintained in 5% L929. To generate immortalized mouse macrophages (iBMDMs), BMDMs were infected with J2 recombinant retrovirus (encoding v-myc and v-raf oncogenes) as described (74). iBMDMs were passaged for 3 to 6 months and were slowly weaned off of L929 conditioned media until they stabilized into cell lines. Human foreskin fibroblasts (HFF, SCRC- 1041, ATCC) were immortalized using a human telomerase-expressing retrovirus (pWZL-Blast-Flag-HA-hTERT, 22396, Addgene).

Bacteria were prepared as previously described (61) with the following exceptions. *B. burgdorferi* was grown to mid-exponential phase, centrifuged at 6600 x g for 8 minutes, washed twice in PBS, and resuspended in DMEM (D5796, Millipore Sigma) with 10% FBS (VWR, 97068-085). Spirochetes were enumerated using dark-field microscopy and diluted to the appropriate MOI. Where indicated, plates were spun following the addition of bacteria to mammalian cells for 5 minutes at 300 x g. Small molecule inhibitors were added to MEFs one hour prior to *B. burgdorferi* infection. cGAS inhibitor RU.521 (HY-114180, MedChemExpress) and STING inhibitor H-151 (HY-112693, MedChemExpress) were added at 10 mM and 0.5 mM, respectively (75, 76). MEFs or BMDMs were transfected with 2 µg/ml Interferon Stimulatory DNA (ISD, tlrl-isdn, InvivoGen) complexed with Lipofectamine 2000 (11668019, ThermoFisher) in Opti-MEM media (11058021, Gibco) for 5-20 minutes (73).

### Quantitative PCR and RT-PCR

RNA was isolated from mammalian cells using the Quick-RNA Micro Prep Kit (R1051; Zymo Research) according to the manufacturer’s instructions. Between 300-500 ng of RNA was standardized across samples from each experiment and converted to cDNA with the qScript cDNA Synthesis Kit (95047, QuantaBio). Quantitative PCR (qPCR) was performed on cDNA using the PerfecTa SYBR Green FastMix (95072, Quantabio) and primers listed in **Table I**. Each biological sample was assayed in triplicate. Relative expression was determined for each triplicate after normalization against a housekeeping gene (*Bactin* or *Gapdh*) using the 2^-ΔΔ*CT*^ method. DNA contamination of RNA samples was evaluated in a single, no RT reaction for each primer set.

### Immunoblotting

Protein was collected from cells lysed in 1% NP-40 buffer (50 mM Tris pH 7.5, 0.15 M NaCl, 1 mM EDTA, 1% NP-40 and 10% glycerol) supplemented with protease inhibitor (04693159001, Roche) and spun for 10 minutes at 17,000 x g at 4°C. The supernatant was collected and stored at -80°C. Protein lysates were quantified using the micro-BCA assay (23235, ThermoFisher Scientific, Waltham, MA). Immunoblotting was performed as described in (77). Briefly, between 20-30 mg protein was run on 10-20% SDS–PAGE gradient gels and transferred onto 0.22 μM PVDF membranes (1620177, Bio-Rad). After air drying to return to a hydrophobic state, membranes were incubated in primary antibodies (**Table II**) at 4 °C overnight in 1X PBS containing 1% casein, HRP-conjugated secondary antibody at room temperature for 1 h, and then developed with Luminata Crescendo Western HRP Substrate (WBLUR0500, Millipore).

### Immunofluorescence microscopy

Cells were seeded on 12- or 18-mm sterile coverslips, allowed to adhere overnight, and infected as described above. At the conclusion of infection, cells were washed with DMEM and then 1X PBS, fixed with 4% paraformaldehyde for 15 minutes at room temperature, and washed twice with 1X PBS for 5 minutes each. Cells were permeabilized with 0.1% Triton X-100 in PBS for 5 minutes at RT, washed twice with 1X PBS and blocked for 30 minutes in PBS containing 5% FBS. Each coverslip was stained with primary (**Table II**) and secondary antibodies for 1 hour each. Cells were washed in 1X PBS with 5% FBS three times after each stain for 5 minutes each. After the last wash, cells were incubated with CellMask Green (H32714, Invitrogen) for 30 minutes at 1:500 dilution in 1X PBS. Then, cells were washed 2 times more for 5 minutes each with 1X PBS. After the last wash, cells were incubated with 4’,6-diamidino-2-phenylindole (DAPI, 62247, ThermoFisher Scientific) for 2 minutes at 1:2000 dilution in 1X PBS. Cells were washed 2 times more for 5 minutes each with 1X PBS, and coverslips were mounted with ProLong Diamond Antifade Mountant (P36961, Invitrogen) and allowed to dry overnight. Images in Figures 2E and 3A were captured with a LSM 780 confocal microscope (Zeiss) with a 63× oil-immersed objective. Z-stack images were processed using Zeiss ZEN 3.3 software. Images in Figure 3F and Supplemental Figure 1A were taken with an ECLIPSE Ti2 microscope (Nikon) with a 60x oil-immersed or 40X dry objective, respectively, and NIS-Elements AR 5.21.02 software. Images in Figure 3F were imported into Huygens Essential software (v21.4.0) and deconvolved using the ‘aggressive’ profile in the Deconvolution Express application. Images in Supplemental Figure 1E were captured with an Olympus FV3000 confocal laser scanning microscope and a 60X oil-immersed objective.

For quantification of *B. burgdorferi* associated with cultured MEF and HFF cells (Supplemental Figure 1A), tiled images from infected cells at 24 hours post infection (hpi) were taken with 40X objective. From each field, the number of cells with more than one *B. burgdorferi* present the cell area, defined by positive CellMask staining, was annotated. *B. burgdorferi* positive cells were divided by the total number of cells per field, defined by counting DAPI positive nuclei, and multiplied by 100 to calculate the percentage of cells with *B. burgdorferi* associated. Four fields for each cell line 6 and 24 hours post infection were quantified from duplicate biological samples, and the average of each cell line across replicates and fields was plotted.

### Mouse infection with *B. burgdorferi* with in vivo bioluminescent imaging

Bioluminescent images of mice were collected as previously described (70, 78). Groups of 5 WT, 5 cGAS^KO^, and 5 STING^KO^ male mice 6-11 weeks old were subcutaneously injected with 100 µl of 10^5^ ML23 pBBE22*luc.* Prior to imaging, 5 mg of D-luciferin (Goldbio, St. Louis, MO) was dissolved in PBS and administered to all except one mouse per group via intraperitoneal injection and anesthetized with isoflurane for imaging. Mice were imaged at 1 hour and 1, 4, 7, 10, 14, 21, and 28 days post-infection (dpi) using the Perkin Elmer IVIS Spectrum live imaging system. Bioluminescence from treated mice were normalized to the untreated mouse from each group. At 28 dpi, inguinal lymph nodes and skin flanks were collected and transferred to BSKII with 6% NRS for outgrowth assays.

### Histopathology

Samples of joints and hearts were collected from each mouse following euthanasia at 28 and 35 dpi, fixed by immersion in 10% neutral buffered formalin at room temperature for 48 hours, and stored in 70% ethanol before embedding in paraffin, sectioning at 5-6 µm, and staining with hematoxylin and eosin by AML Laboratories, Inc. (Jacksonville, Florida). The tissue sections were examined using brightfield microscopy in a blinded manner by a board-certified anatomic veterinary pathologist and ordinally scored for the degree of mononuclear infiltration. Score of 1 represented minimal infiltration (<5 cells/400X field) and increased to 4 for abundant infiltration (>30cells/400X field). Tissue sections were also scored for distribution of mononuclear inflammatory cells as focal (1), multifocal (2), or diffuse (3).

### Statistical analyses

Statistical analysis was performed in GraphPad Prism (GraphPad Software, Inc., La Jolla, CA). Statistical significance was determined by *p* ≤ 0.05. Specific tests are detailed in the figure legends. Error bars in figures represent standard error of the mean (SEM) based on the combined triplicate biological samples for all cell culture studies. All cell culture-based results (Figures 1-3, Supplemental Figures 1-2) are representative of at least three independent experiments. Four biological replicates were utilized in in vivo bioluminescence imaging (Figure 4) to determine statistical significance.

**Figure 1:**
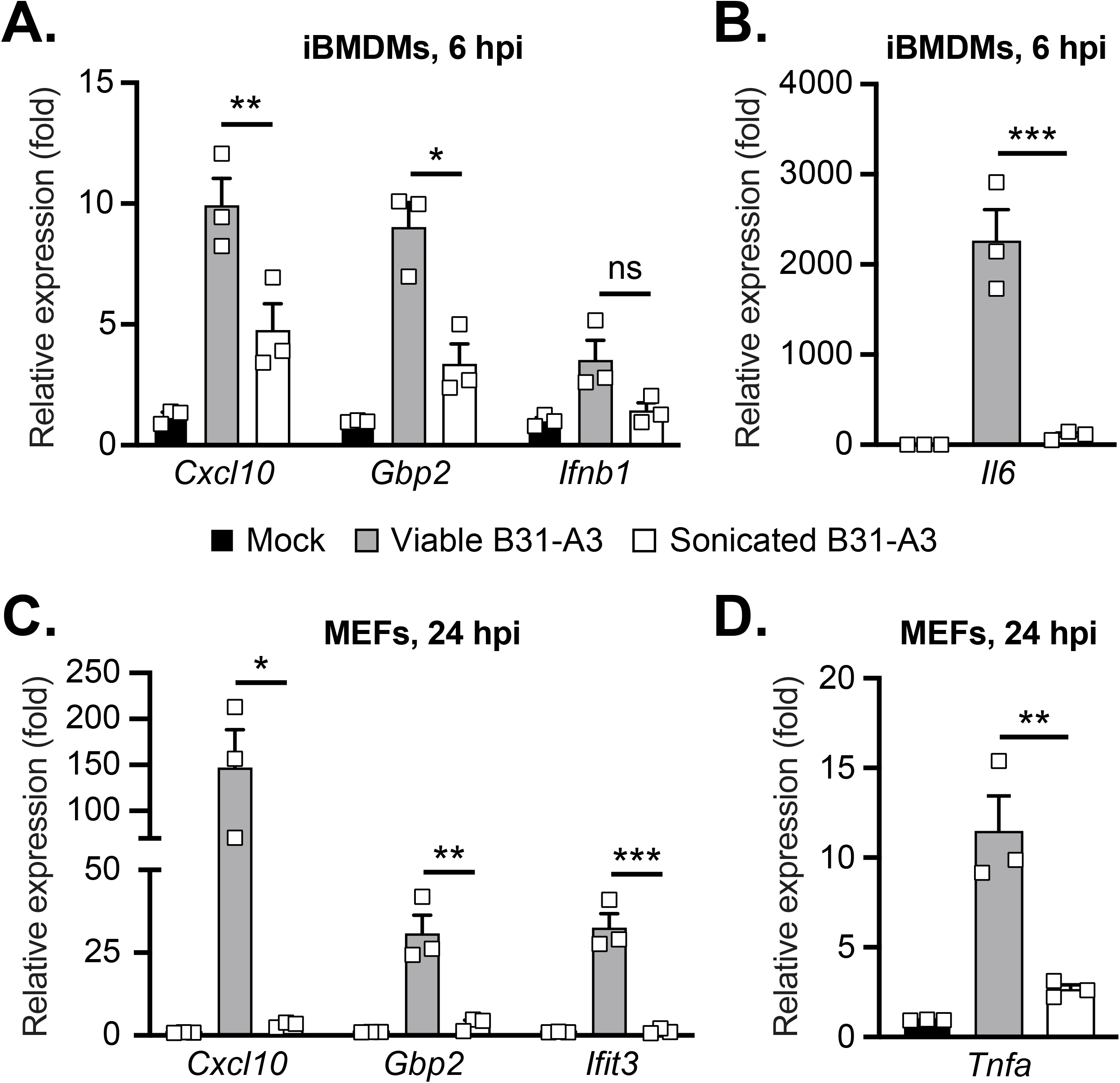
Viable *B. burgdorferi* induce robust proinflammatory and type I interferon responses in mouse macrophages and fibroblasts. Immortalized mouse bone marrow-derived macrophages (iBMDMs) or murine embryonic fibroblasts (MEFs) were co-cultured with viable or sonicated *B. burgdorferi* strain B31-A3 at a MOI of 20. (**A**) Fold changes in iBMDM transcripts encoding interferon-stimulated genes (ISGs) (*Cxcl10* and *Gbp2*), *Ifnb1*, or (**B**) the pro-inflammatory cytokine *Il6* were analyzed by qRT-PCR after 6 hours of co-culture. (**C**) Fold changes in MEF transcripts encoding ISGs (*Cxcl10, Gbp2* and *Ifit3*) and the (**D**) inflammatory cytokine *Tnfa* were analyzed by qRT-PCR after 24 hours of co-culture. Error bars represent ± SEM of biological triplicate samples. Significance was determined by one-way ANOVA Tukey post hoc for all panels. *** *p* < 0.001, \*\**p* < 0.01, * *p* < 0.05, ns, not significant.

## RESULTS

### Viable *B. burgdorferi* elicits a robust type I interferon response in mouse macrophages and fibroblasts

We first evaluated the ability of immortalized bone marrow-derived macrophages (iBMDMs) and embryonic fibroblasts (MEFs) derived from C57BL/6J mice to upregulate interferon stimulated gene (ISG) and inflammatory cytokine transcripts when co-incubated with viable or sonicated *B. burgdorferi* B31-A3 at a MOI 20 (**Figure 1**). Consistent with earlier reports (19, 60), both viable and sonicated B31-A3 were able to induce interferon beta (*Ifnb1*) and ISGs (C-X-C motif chemokine ligand 10 (*Cxcl10*) and guanylate-binding protein 2 (*Gbp2*)), as well as pro-inflammatory cytokine interleukin-6 (*Il6*) expression, in iBMDMs (**Figure 1A-B**). Viable *B. burgdorferi* more potently induced ISGs and *Il6*, inducing expression levels 4-10 fold higher than sonicated bacteria. Likewise, we found that *B. burgdorferi* elicited ISGs and tumor necrosis factor alpha (*Tnfa*) in primary MEFs, with live spirochetes triggering more robust responses compared to sonicated bacteria (**Figure 1C-D**). Collectively, these data indicate that live *B. burgdorferi* engages robust IFN-I-associated ISG expression in both BMDMs and non-phagocytic MEFs, which lack a full repertoire of TLRs and other pattern recognition receptors (41).

### *B. burgdorferi* engages the cGAS-STING pathway to induce type I interferon responses in murine macrophages

Lipoprotein rich *B. burgdorferi* predominately engage TLR2, but can also trigger other TLRs to induce inflammatory cytokines and interferons (18, 19, 24, 36). However, a role for cytosolic DNA sensing in the innate immune response to *B. burgdorferi* remains unknown. To examine this directly, we co-cultured primary BMDMs from wild-type (WT), cGAS^KO^, or STING^KO^ mice on a C57BL/6J background with *B. burgdorferi* B31-A3 and assessed ISG and cytokine transcript induction by qRT-PCR (**Figure 2**). We first confirmed that cGAS^KO^ BMDMs were hyporesponsive to immunostimulatory DNA (ISD) delivered into the cytosol by transfection (73). As expected, the induction of ISGs *Cxcl10* and *Gbp2* in cGAS^KO^ BMDMs was significantly reduced relative to WT macrophages (**Figure 2A**). After 6 hours of co-culture with B31-A3, the expression of ISGs *Cxcl10* and *Gbp2*, as well as *Ifnb1* transcripts, were significantly reduced in cGAS^KO^ BMDMs compared to WT (**Figure 2B**). In contrast, the levels of *Il-6* and *Tnfa* transcripts were similar in both WT and cGAS^-/-^ BMDMs (**Figure 2C**). These results indicate that the cGAS pathway is not a robust inducer of pro-inflammatory genes in BMDMs exposed to *B. burgdorferi and* are consistent with the notion that TLR2 or other TLRs sense *B. burgdorferi* ligands to engage nuclear-factor kappa-B (NF-κB)-dependent cytokines. Similar results were obtained from protein analysis of cGAS^KO^ and STING^KO^ macrophages 9 hours after incubation with B31-A3, with notable reductions in ISGs interferon-induced protein with tetratricopeptide repeats 1 (IFIT1) and Z-DNA binding protein 1 (ZBP1) in both cGAS- and STING-deficient BMDMs (**Figure 2D**). Finally, confocal imaging revealed that *B. burgdorferi* are internalized by BMDMs after a 3 hour incubation (**Figure 2E**). Antibody staining against the Outer Surface Protein A (OspA) of *B. burgdorferi* showed coiled and degraded DAPI positive spirochetes in the macrophage cytoplasm, as well as punctate, DAPI negative OspA staining, indicative of bacterial destruction. *B. burgdorferi* OspA also co-localized with the cytosolic autophagy marker p62/Sequestosome-1 (SQSTM1), suggesting spirochete degradation via macrophage autophagy and/or LC3-associated phagocytosis pathways (79–81). Collectively, these data indicate that *B. burgdorferi* are internalized by murine macrophages and trigger the cGAS-STING-IFN-I signaling axis.

**Figure 2:**
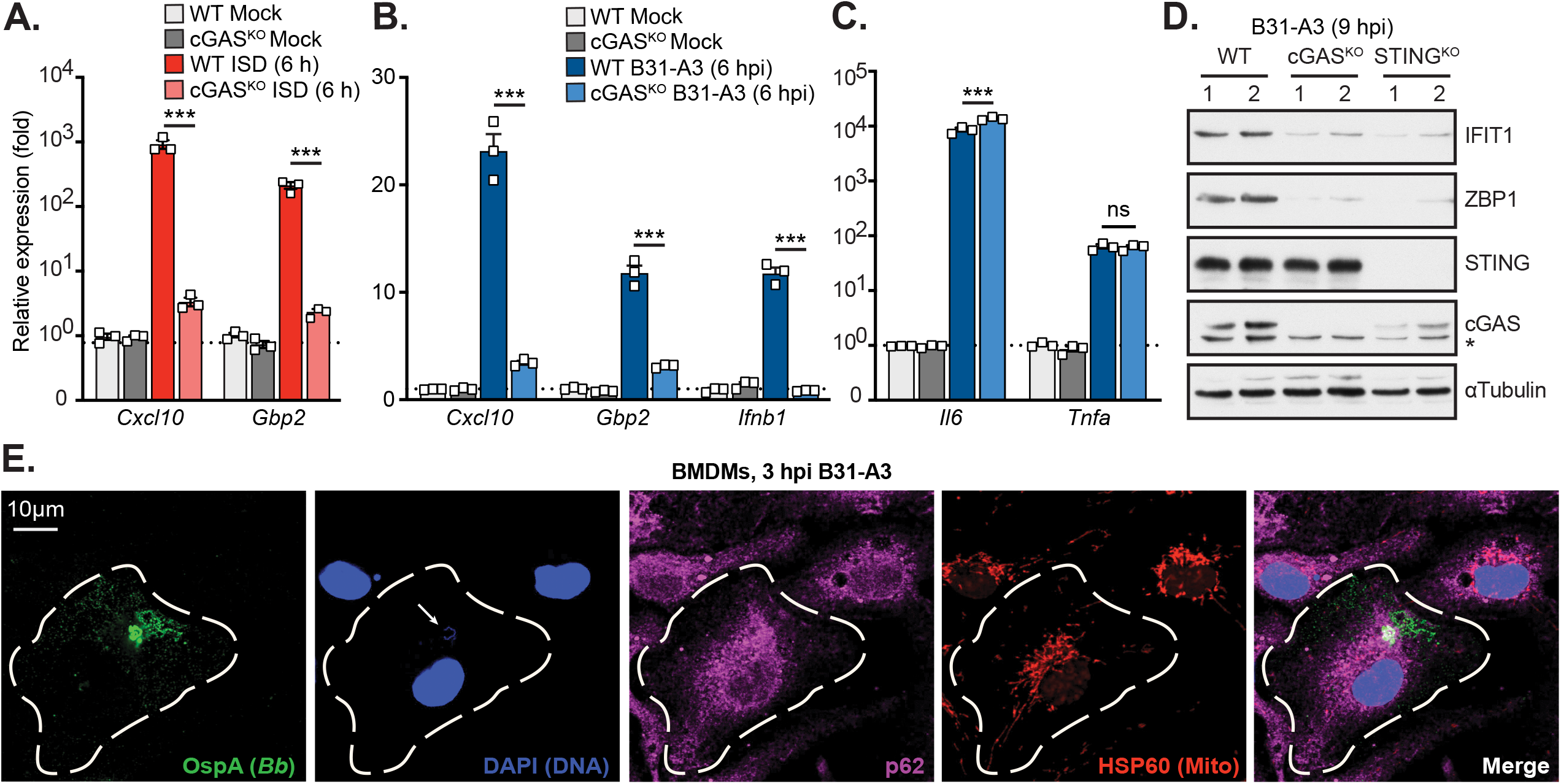
cGAS is required for robust ISG expression in mouse macrophages exposed to live *B. burgdorferi*. (**A**) qRT-PCR of ISG transcripts (*Cxcl10, Gbp2*) from wild-type (WT) and cGAS knockout (cGAS^KO^) bone marrow-derived macrophages (BMDMs) 6 hours after transfection with immunostimulatory DNA (ISD) or mock transfected. (**B-C**) qRT-PCR of transcripts encoding ISGs (*Cxcl10, Gbp2*) and *Ifnb1* (**B**) or inflammatory cytokines (*Il6* and *Tnfa*) (**C**) in WT and cGAS^KO^ BMDMs exposed to live *B. burgdorferi* strain B31-A3 for 6 hours at MOI 20. Fold expression values in **A**, **B**, **C** are plotted relative to WT mock samples. (**D**) Western blots of WT, cGAS^KO^, and STING deficient (STING^KO^) BMDMs mock infected or exposed to B31-A3 at a MOI 20 for 9 hours. Each lane represents a biological duplicate representative of three independent experiments. Non-specific band indicated by *. (**E**) WT BMDMs were co-cultured with strain B31-A3 at MOI 20 for 3 hours on coverslips. Cells were fixed and stained with antibodies against Borrelial outer surface protein A (OspA), autophagy marker p62, mitochondrial heat shock protein 60 (HSP60), counterstained with DAPI, and subjected to confocal microscopy. Arrow indicates *B. burgdorferi* DNA stained with DAPI. Error bars in **A**, **B**, **C** represent ± SEM of triplicate biological samples. One-way ANOVA Tukey post hoc was used in **A**, **B**, **C** to determine significance. \*\*\**p* < 0.001, ns, not significant.

### *B. burgdorferi* engages the cGAS-STING pathway in fibroblasts

To broaden our findings beyond murine macrophages, we exposed MEFs and telomerase immortalized human foreskin fibroblasts (HFF) to *B. burgdorferi* B31-A3. We observed that OspA positive *B. burgdorferi* associated with approximately 15% of MEFs (**Supplemental Figure 1A-B**) and 33% of HFFs (**Supplemental Figure 1C-D**) after co-culture, consistent with a prior study (54). Confocal immunofluorescent imaging with Z-stack reconstitution revealed cell-associated *B. burgdorferi* in the same focal plane with mitochondria (**Figure 3A**), suggestive of spirochete internalization. Additional imaging analysis revealed that coiled and degraded spirochetes strongly co-localized with the intracellular autophagy marker p62, further documenting that *B. burgdorferi* can access the fibroblast cytoplasm (**Supplemental Figure 1E**). Similar to our results in MEFs, exposure of HFFs to live *B. burgdorferi* increased expression levels of ISGs (*IFI44L*, *IFNB1*) and the pro-inflammatory cytokine gene *TNFA* (**Figure 3B**). Although the synthetic TLR2 ligand PAM3CSK4 (Pam3) induced *TNFA* levels similar to live B31-A3, we did not observe ISG induction, suggesting that *B. burgdorferi* lipoprotein engagement of TLR2 is not responsible for IFN-I responses in HFFs. Taken together, these results demonstrate that *B. burgdorferi* association and internalization within a minority of cultured cells is sufficient to induce robust IFN-I responses.

**Figure 3:**
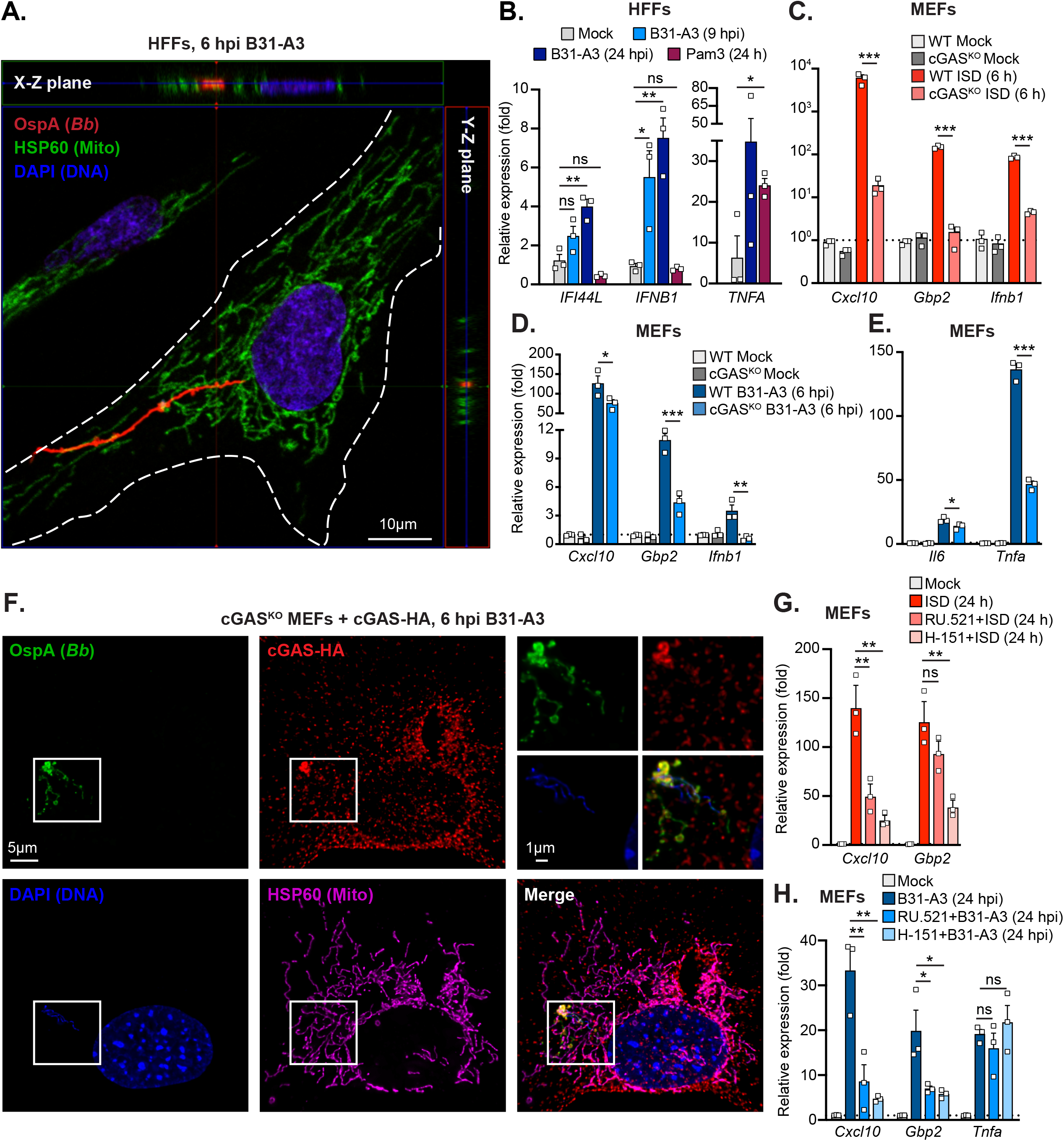
cGAS governs ISG expression in fibroblasts exposed to live *B. burgdorferi*. (**A-B**) Telomerase immortalized human foreskin fibroblasts (HFFs) were co-cultured with *B. burgdorferi* at a MOI of 20. (**A**) After 6 hours of incubation, cells were fixed and stained with antibodies against Borrelial outer surface protein A (OspA) and mitochondrial heat shock protein 60 (HSP60), counterstained with DAPI, and subjected to confocal microscopy. Z-stack image reconstitution was performed to localize internalized *B. burgdorferi* with mitochondria. (**B**) After 9 and 24 hours of co-culture with B31-A3 or stimulation with the Toll-like receptor ligand PAM3CSK4, RNA was extracted for qRT-PCR analysis of ISG (*IFI44L* and *IFNB1*) and inflammatory cytokine (*TNFA*) transcripts. (**C**) qRT-PCR of ISG (*Cxcl10, Gbp2*) and *Ifnb1* transcripts from wild-type (WT) and cGAS knockout (cGAS^KO^) murine embryonic fibroblasts (MEFs) 6 hours after transfection with immunostimulatory DNA (ISD) or mock transfected. (**D- E**) qRT-PCR of transcripts encoding ISGs (*Cxcl10, Gbp2*) and *Ifnb1* (**D**) or inflammatory cytokines (*Il6* and *Tnfa*) (**E**) in WT and cGAS^KO^ MEFs exposed to live *B. burgdorferi* strain B31-A3 for 6 hours at MOI 20. (**F**) Immortalized cGAS^KO^ MEFs stably reconstituted with HA- tagged cGAS were co-cultured with strain B31-A3 at MOI 20 for 6 hours on coverslips. Cells were fixed and stained with antibodies against Borrelial outer surface protein A (OspA), HA (cGAS-HA), and mitochondrial heat shock protein 60 (HSP60), then counterstained with DAPI, and subjected to fluorescence microscopy. (**G-H**) WT MEFs were treated with the cGAS inhibitor RU.512 or the STING inhibitor H-151 for one hour prior to ISD transfection (**G**) or addition of *B. burgdorferi* B31-A3 (**H**). After 24 hours of incubation, qRT-PCR analysis of ISG (*Cxcl10* and *Gbp2*) or inflammatory cytokine (*Tnfa*) transcripts was run. Fold expression values in **B-E** and **G**-**H** are plotted relative to WT mock samples, and error bars represent ± SEM of triplicate biological samples. One-way ANOVA Tukey post hoc was used to determine significance. \*\*\**p* < 0.001, \*\**p* < 0.01, \**p* < 0.05, ns, not significant.

To next examine whether cGAS contributes to IFN-I responses in fibroblasts exposed to *B. burgdorferi*, primary WT and cGAS^KO^ MEFs were co-cultured with B31-A3 and subjected to qRT-PCR analysis for ISG and pro-inflammatory transcripts. We first confirmed that cGAS^KO^ MEFs were hyporesponsive to immunostimulatory DNA (ISD) delivered into the cytosol by transfection (**Figure 3C**). After co-culture with *B. burgdorferi*, we noted that the induction of ISG (*Cxcl10* and *Gbp2*) and *Ifnb1* transcripts was significantly reduced or entirely abrogated in cGAS^KO^ MEFs relative to WT controls (**Figure 3D**). Although *Il6* transcripts were similar between *B. burgdorferi* exposed WT and cGAS^KO^ MEFs, *Tnfa* expression was significantly reduced in the absence of cGAS (**Figure 3E**). Immunofluorescence microscopy revealed that *B. burgdorferi* associated with MEFs after a 6 hour incubation, similar to results in BMDMs and HFFs (**Figure 3F**). Antibody staining against the OspA lipoprotein revealed intact, DAPI positive spirochetes in the MEF cytoplasm, as well as a punctate OspA foci that were DAPI negative. Consistent with a role for cGAS in sensing internalized *B. burgdorferi* DNA, we observed co-localization of HA-tagged cGAS with coiled spirochetes staining positive for both OspA and DAPI (**Figure 3F**). To further document a requirement for the cGAS-STING pathway in the IFN-I response to *B. burgdorferi*, we employed RU.521, a specific cGAS inhibitor, and H- 151, a specific STING inhibitor, to block the pathway during co-culture. Both inhibitors were effective at reducing ISG transcripts (*Cxcl10* and *Gbp2*) induced by transfection of ISD (**Figure 3G**). MEFs exposed to cGAS and STING inhibitors exhibited reduced ISG expression relative to vehicle treated MEFs, with no effects on *Tnfa* induction (**Figure 3H**). Taken together, these data indicate that *B. burgdorferi* associates with fibroblasts in culture, leading to the cGAS-mediated sensing of borrelial DNA from lysed or damaged spirochetes. Additional experiments in STING^KO^ MEFs revealed markedly reduced *Gbp2* and *Ifnb1* expression, but little change in pro-inflammatory cytokine transcripts, after challenge with *B. burgdorferi* (**Supplemental Figure 2 A-B**). MEFs deficient in the type I interferon receptor (IFNAR) also exhibited impaired expression of the ISG *Gbp2*, but not cytokine transcripts, suggesting that ISG induction in MEFs is dependent on autocrine and/or paracrine signaling via IFNAR. In contrast, *B. burgdorferi* infection of Mitochondrial Antiviral Signaling protein null MEFs (MAVS^KO^), which cannot signal in response to intracellular dsRNA, triggered levels of *Gbp2* and *Ifnb1* mirroring WT cells (**Supplemental Figure 2A**). This indicates that MEFs co-cultured with *B. burgdorferi* respond to DNA, not RNA, ligands to induce IFN-I. Expression of pro-inflammatory cytokine transcripts *Il6* and *Tnfa* were unchanged or modestly altered in STING^KO^, MAVS^KO^, and IFNAR^KO^ MEFs, demonstrating all MEF lines remained responsive to *B. burgdorferi* lipoprotein engagement of TLR2 during co-incubation (**Supplemental Figure 2B**). Moreover, qRT-PCR to detect internal levels of borrelial flagellar gene, *flaB,* within host cells revealed roughly equivalent levels of *flaB* among WT and mutant MEF lines (**Supplemental Figure 2C**). This indicates that altered ISG expression in STING and IFNAR deficient cells is not due to changes in the ability of viable *B. burgdorferi* to associate with these MEFs.

### cGAS-STING modulates inflammation during *B. burgdorferi* infection in vivo

To determine roles for the cGAS-STING pathway in borrelial dissemination, tissue colonization, and inflammation during mammalian infection, bioluminescent *B. burgdorferi*, ML23 pBBE22*luc*, was monitored in real time by in vivo imaging in C57BL/6 WT, cGAS^KO^, and STING^KO^ mice (**Figure 4**). Mice were injected with D-luciferin prior to imaging and one mouse in each group was not injected to serve as a bioluminescence background control (**Figure 4A**). The absence of cGAS or STING did not alter the kinetic dissemination of *B. burgdorferi* or the borrelial load as observed in the images and by quantitative analysis of bioluminescence emission (**Figure 4A-B**). Outgrowth from infected tissues confirmed that all genotypes were colonized with viable *B. burgdorferi* 28 days post infection (**Figure 4C**). To investigate whether the cGAS-STING pathway impacts the development of arthritic phenotypes, tibotarsal joints from WT and cGAS^KO^ mice were collected for histopathology. Consistent with prior reports (82), *B. burgdorferi* induced mild arthritis in WT C57BL/6J mice at 28 days post infection (**Supplemental Figure 3A**). Interestingly, H&E staining revealed that the joints of cGAS^KO^ mice exhibited reduced synovial papillary hyperplasia, immune cell infiltration, and overall joint pathology scores, suggesting a trend toward reduced inflammation the 28 day timepoint. These results suggest that the cGAS-STING pathway may contribute to the development of inflammation during mammalian infection, without impacting the ability of *B. burgdorferi* to readily disseminate and colonize secondary tissues.

**Figure 4:**
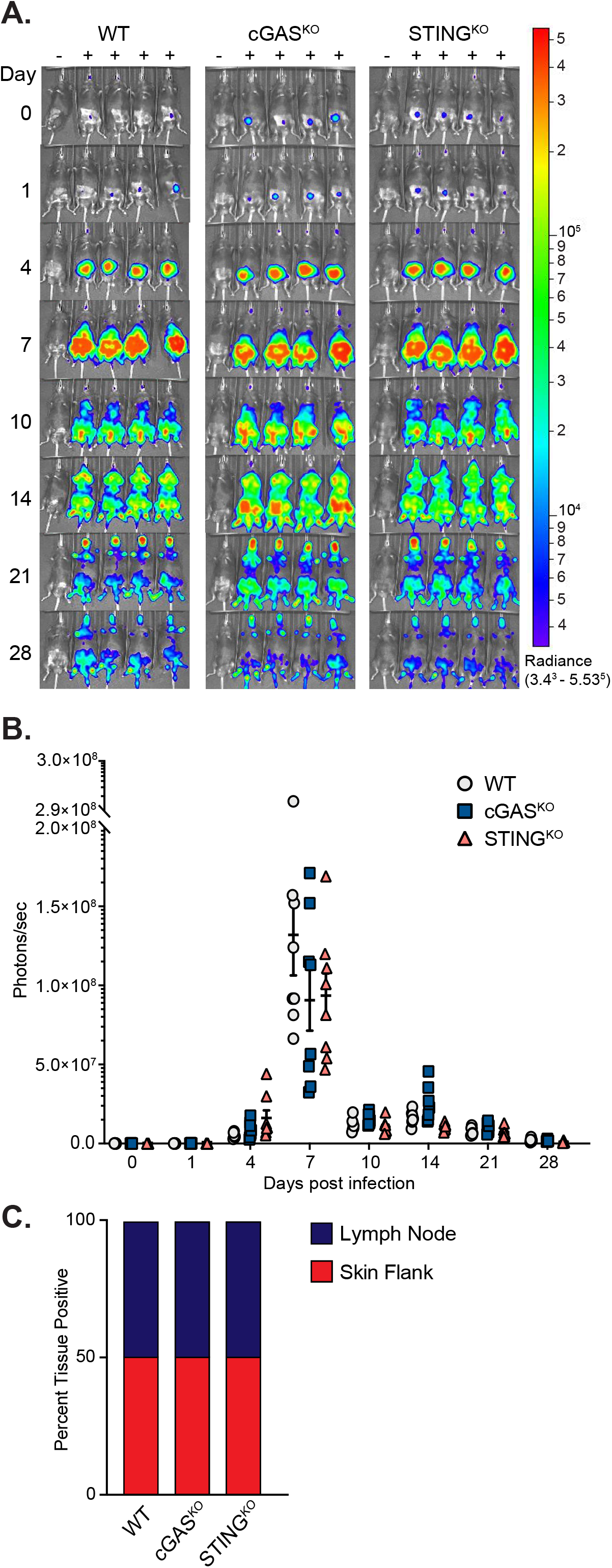
*B. burgdorferi* infection kinetics and tissue colonization are similar among WT, cGAS, and STING deficient mice. WT, cGAS^KO^, and STING^KO^ mice on the C57BL/6J background were infected with 10^5^ ML23 pBBE22*luc B. burgdorferi (n = 10/strain, 30 mice total)*. (**A**) Mice were selected randomly and treated with D-luciferin at 1, 4, 7, 10, 14, 21, and 28 days post infection for *in vivo* imaging. For each mouse genotype, the mouse in the first column did not receive D-luciferin to serve as a background control. All images were normalized to the 3.4×10^3^ to 5.53×10^5^ radiance (p/s/cm^2^/sr) and displayed on the same color spectrum scale (right). (**B**) Quantification of bioluminescence in p/s. (**C**) Qualitative assessment of dissemination through tissue outgrowth. At days 28, 5 mice per strain were sacrificed after completion of imaging. An inguinal lymph node and skin flank were collected for *in vitro* cultivation. Percent of positive cultures is indicated by the y-axis. There was no significant difference in bioluminescence or tissue outgrowth across all mouse genotypes.

## DISCUSSION

*B. burgdorferi* elicits robust innate and adaptive immune responses that involve induction of both IFN-I (IFN-α and β) and IFN-II (IFN-γ) cytokines that drive expression of an overlapping family of interferon-stimulated genes (ISGs) (12, 31, 35, 83, 84). Synovial tissue from patients with post-infection Lyme arthritis expressed ISGs that are associated with both type I and type II interferons (25, 35, 35, 85). Antibody-mediated IFNAR blockade results in a significant reduction in joint inflammation in mice (12), and the inhibition of IFN-γ delays sustained ISG expression and inflammation during *B. burgdorferi* infection (12, 86). A number of prior studies have focused on identifying the signaling mechanisms responsible for IFN-I and ISGs elicited by *B. burgdorferi*. Both in vitro and in vivo studies have demonstrated that TLR adaptors MyD88 and TRIF are not the primary contributors to IFN-I induction (12, 13, 22, 87–90). Specifically, in the absence of TLR2 or TLR5, innate immune receptors that sense lipoproteins or flagella of degraded *B. burgdorferi,* respectively, the induction of ISGs and joint swelling is similar to WT mice (12, 87–89). Studies using human cells have implicated nucleic acid sensing, endosomal localized TLRs (TLR 7, 8, and 9) as regulators of IFN-I induction in human peripheral blood mononuclear cells (PBMCs) challenged with *B. burgdorferi* ex vivo (13, 18, 19, 90). However, TLR9 inhibition does not impair ISG induction in BMDMs exposed to *B. burgdorferi*, in contrast to PBMCs (19, 22, 85). Thus, it is likely that multiple innate immune pathways are responsible for IFN-I induction during *B. burgdorferi* infection. Differences in IFN-I and inflammatory responses across independent studies might also be explained by the use of distinct borrelial strains that vary in invasion and inflammation (11, 91, 92).

Fibroblast and endothelial cells, which do not express a full complement of TLRs, are necessary for *B. burgdorferi* induced IFN-I responses in joints (25, 41). Thus, TLR detection of *B. burgdorferi* nucleic acids is likely not the predominant pathway governing IFN-I responses during *B. burgdorferi* infection in the mammalian host. Therefore, we hypothesized that *B. burgdorferi* might engage an intracellular innate immune pathway leading to IFN-I. Using primary fibroblasts and macrophages from a panel of C57BL/6J mice lacking intracellular nucleic acid sensors or downstream adaptors, we find that the DNA sensing, cGAS-STING pathway is critically required for robust induction of ISGs after exposure to live *B. burgdorferi*.

This finding was confirmed through small molecule inhibitor studies, as we observed reduced ISG responses to *B. burgdorferi* when cGAS or STING were inhibited in MEFs. In contrast, we show that deficiency of the main RIG-I-like receptor adaptor MAVS does not impact expression of ISGs or pro-inflammatory cytokine transcripts in fibroblasts, indicating that intracellular sensing of *B. burgdorferi* RNA is not a major contributor to IFN-I responses in cultured fibroblasts.

The phenomenon of host cell internalization by *B. burgdorferi* has not been observed during experimental infection in vivo; however, prior studies have shown that *B. burgdorferi* can become internalized into both phagocytic (macrophages, monocytes, and dendritic cells) and non-phagocytic (endothelial, fibroblast, and neuroglial cells) cells in culture (54, 57–62, 93–95).

Our immunofluorescence microscopy analyses revealed borrelial OspA staining associated with a proportion of cultured mouse and human fibroblasts after short incubations with *B. burgdorferi* B31-A3. In both macrophages and fibroblasts, we observed OspA colocalization with the intracellular autophagy adaptor p62, which substantiates the notion that spirochetes interact with mammalian cells and can become internalized into the cytoplasm. Recent work has revealed that the autophagy machinery regulates the cGAS-STING pathway, as p62 is found in close proximity to TBK1, a key kinase necessary for IFN-I induction by STING (96). In addition, phosphorylation of p62 by TBK1 is important for the negative regulation and turnover of STING (97). Activation of cGAS by the obligate intracellular pathogen *Mycobacterium tuberculosis* can drive selective autophagy for clearance of intracellular bacteria (45). Interestingly, we observed coiled and degraded *B. burgdorferi* in BMDMs and MEFs colocalizing with p62, suggesting that the autophagy could be a mechanism to turnover internalized spirochetes. Autophagy has also been linked to inflammatory cytokine responses during *B. burgdorferi* infection in vitro (79). Therefore, internalization of *B. burgdorferi* and autophagic targeting may serve to fully engage intracellular signaling leading IFN-I or other innate immune responses.

The cGAS-STING pathway is a major inducer of IFN-I in response to both exogenous, pathogen-derived DNA, as well as host mitochondrial and nuclear DNA (45, 51, 52, 72, 98–100). cGAS non-discriminately binds double stranded DNA and produces cGAMP that binds and activates STING on the ER. This leads to recruitment and phosphorylation of the kinase TBK1, which phosphorylates and induces nuclear translocation of transcription factors IRF3 or 7 (101). Miller et al demonstrated that *B. burgdorferi* engages IFN-I response genes in an IRF3- dependent manner, further supporting that the signaling machinery downstream of cGAS-STING and other intracellular nucleic acid sensors is required for ISG induction (12). Multiple intracellular bacterial pathogens, including *Mycobacterium tuberculosis, Listeria monocytogenes,* and *Chlamydia trachomatis*, engage the cGAS-STING axis to induce IFN-I and influence pathogenesis (45, 50, 102–105). More recently, extracellular bacterial pathogens *Pseudomonas aeruginosa* and Group B *Streptococcus* (GBS) have been shown to trigger IFN-I responses through cGAS-STING in macrophages and dendritic cells (52, 106). The mechanisms by which *B. burgdorferi* and other extracellular bacteria engage cGAS remain unclear. However, we have observed recruitment and co-localization of cGAS with intracellular borrelia in MEFs, suggesting that internalized spirochetes may shed DNA or undergo host-induced membrane damage that liberates bacterial genomic material for detection by cGAS. Bacterial pathogens can also cause host cell stress, resulting in the release and accumulation of nuclear or mitochondrial DNA (mtDNA) (107, 108). Mitochondrial stress and mtDNA release, in particular, is linked to innate immune responses through the cGAS-STING axis and is associated with autoimmune disorders (i.e. Lupus and rheumatoid arthritis) and other conditions characterized by elevated IFN-I (98). Thus, it is possible that the IFN-I response induced by *B. burgdorferi* is at least partially dependent on host mtDNA or nuclear DNA sensing. It is likely that *B. burgdorferi* engages distinct mechanisms in phagocytic and non-phagocytic cells to elicit IFN-I responses via the cGAS-STING pathway, and future work is required to reveal the molecular mechanisms involved.

An IFN-I response occurs early in mammalian infection resulting in the development of inflammation and arthritis that persist in part due to IFN-γ during later stages of disease (32, 35, 83, 84). We therefore investigated how absence of cGAS and STING impacted the kinetics of borrelial infection in mice using in vivo imaging to track bioluminescent *B. burgdorferi* in real time through the progression of disease (70, 109). Bioluminescent *B. burgdorferi* was able to successfully disseminate and colonize secondary tissues independently of cGAS or STING. Joint inflammation also showed a moderate, although not statistically significant, reduction at 28 dpi in cGAS^KO^ mice, suggesting that cGAS-STING signaling to IFN-I or other inflammatory response pathways might contribute to arthritic phenotypes during *B. burgdorferi* infection. It is important to note that the absence of cGAS did not eliminate inflammation entirely, indicating that *B. burgdorferi* associated arthritis is multifactorial and involves additional innate and adaptive immune pathways as have been reported by others (13, 17, 19, 20, 90).

While this study is impactful in that it is the first to link the cGAS-STING pathway to *B. burgdorferi* induced IFN-I responses, it is not without limitations. C3H is the preferred model for borrelial infection because this background develops inflammation and pathologic disease at a significantly higher level than C57BL/6 or Balb/c mice (34, 82). An elegant study identified the *Bbaa1* locus in C3H mice responsible for IFN-I responses observed during borrelial infection and showed that introduction of this locus into C57BL/6 mice yielded similar inflammatory responses to C3H (29). Our study employed WT, cGAS, and STING knockout strains on a pure C57BL/6 background as they are commercially available and well-characterized. We employed highly invasive and inflammatory RST1 *B. burgdorferi* B31-A3 for our studies and were able to observe IFN-I induction even in cells from the C57BL/6 background. These responses were markedly attenuated or lost when using sonicated *B. burgdorferi* B31-A3. Other studies have revealed weak or absent ISG expression during in vitro challenge with *B. burgdorferi*, although most utilized sonicated bacteria or less inflammatory RST3 strains (12, 22, 82, 87). This indicates that the C57BL/6 background is not devoid of IFN-I responsiveness to borrelial infection. Our work and previous studies show C57BL/6 mice and cell lines do produce measurable IFN-I and inflammatory responses to borrelial infection, and therefore we propose it is an appropriate model for this initial characterization of cGAS-STING in the innate immune response to *B. burgdorferi* (12). However, the development of cGAS or STING knockout lines on the C3H background will be necessary to fully characterize the role of this innate immune pathway in *B. burgdorferi* infection dynamics, tissue-specific inflammation, and arthritic phenotypes in vivo.

In conclusion, our study is the first to show that *B. burgdorferi* triggers the intracellular cGAS-STING DNA sensing pathway to shape IFN-I responses in cultured cells. Future studies are needed to investigate the how borrelial cells initiate IFN-I responses through this pathway by characterizing the source of DNA that binds to cGAS (i.e., bacterial or host) and determining whether borrelial internalization is required. Loss of cGAS or STING does not appear to alter *B. burgdorferi* dissemination or its ability to colonize secondary tissues, but studies in C3H mice lacking this innate immune pathway may reveal differential bacterial kinetics and/or inflammatory responses that do impact the course of infection. Additional work to clarify these open questions may support the development cGAS- and/or STING-based immunotherapeutics that may be effective against active infection or persistent symptoms of *B. burgdorferi*, such as Lyme arthritis.

## ACKNOWLEDGEMENTS

We thank members of the Hyde and West labs for helpful discussions. We thank Dr. Jon Skare for providing the OspA antibody.

## REFERENCES

1. Steere, A. C., F. Strle, G. P. Wormser, L. T. Hu, J. A. Branda, J. W. R. Hovius, X. Li, and P. S. Mead. 2016. Lyme borreliosis. Nat. Rev. Dis. Primer 2: 16090.

2. Stanek, G., and F. Strle. 2018. Lyme borreliosis–from tick bite to diagnosis and treatment. FEMS Microbiol. Rev. 42: 233–258.

3. Radolf, J. D., M. J. Caimano, B. Stevenson, and L. T. Hu. 2012. Of ticks, mice and men: understanding the dual-host lifestyle of Lyme disease spirochaetes. Nat. Rev. Microbiol. 10: 87– 99.

4. Shapiro, E. D. 2014. Lyme disease. N. Engl. J. Med. 371: 684.

5. Kugeler, K. J., A. M. Schwartz, M. J. Delorey, P. S. Mead, and A. F. Hinckley. Estimating the Frequency of Lyme Disease Diagnoses, United States, 2010–2018 - Volume 27, Number 2— February 2021 - Emerging Infectious Diseases journal - CDC..

6. Mead, P. S. 2015. Epidemiology of Lyme disease. Infect. Dis. Clin. North Am. 29: 187–210.

7. Schwartz, A. M. 2017. Surveillance for Lyme Disease — United States, 2008–2015. MMWR Surveill. Summ. 66.

8. Radolf, J. D., K. Strle, J. E. Lemieux, and F. Strle. 2021. Lyme Disease in Humans. Curr. Issues Mol. Biol. 42: 333–384.

9. Bamm, V. V., J. T. Ko, I. L. Mainprize, V. P. Sanderson, and M. K. B. Wills. 2019. Lyme Disease Frontiers: Reconciling Borrelia Biology and Clinical Conundrums. Pathog. Basel Switz. 8: E299.

10. Khatchikian, C. E., R. B. Nadelman, J. Nowakowski, I. Schwartz, M. Z. Levy, D. Brisson, and G. P. Wormser. 2015. Public health impact of strain specific immunity to Borrelia burgdorferi. BMC Infect. Dis. 15: 472.

11. Strle, K., K. L. Jones, E. E. Drouin, X. Li, and A. C. Steere. 2011. Borrelia burgdorferi RST1 (OspC type A) genotype is associated with greater inflammation and more severe Lyme disease. Am. J. Pathol. 178: 2726–2739.

12. Miller, J. C., Y. Ma, J. Bian, K. C. F. Sheehan, J. F. Zachary, J. H. Weis, R. D. Schreiber, and J. J. Weis. 2008. A critical role for type I IFN in arthritis development following Borrelia burgdorferi infection of mice. J. Immunol. Baltim. Md 1950 181: 8492–8503.

13. Cervantes, J. L., S. M. Dunham-Ems, C. J. La Vake, M. M. Petzke, B. Sahay, T. J. Sellati, J. D. Radolf, and J. C. Salazar. 2011. Phagosomal signaling by Borrelia burgdorferi in human monocytes involves Toll-like receptor (TLR) 2 and TLR8 cooperativity and TLR8-mediated induction of IFN-beta. Proc. Natl. Acad. Sci. U. S. A. 108: 3683–3688.

14. Wooten, R. M., M. A. Ying, R. A. Yoder, J. P. Brown, J. H. Weis, J. F. Zachary, C. J. Kirschning, and J. J. Weis. 2002. Toll-like receptor 2 plays a pivotal role in host defense and inflammatory response to Borrelia burgdorferi. Vector Borne Zoonotic Dis 2: 275–8.

15. Marre, M. L., T. Petnicki-Ocwieja, A. S. DeFrancesco, C. T. Darcy, and L. T. Hu. 2010. Human integrin α(3)β(1) regulates TLR2 recognition of lipopeptides from endosomal compartments. PloS One 5: e12871.

16. Dennis, V. A., S. Dixit, S. M. O’Brien, X. Alvarez, B. Pahar, and M. T. Philipp. 2009. Live Borrelia burgdorferi spirochetes elicit inflammatory mediators from human monocytes via the Toll-like receptor signaling pathway. Infect. Immun. 77: 1238–1245.

17. Shin, O. S., R. R. Isberg, S. Akira, S. Uematsu, A. K. Behera, and L. T. Hu. 2008. Distinct roles for MyD88 and Toll-like receptors 2, 5, and 9 in phagocytosis of Borrelia burgdorferi and cytokine induction. Infect. Immun. 76: 2341–2351.

18. Cervantes, J. L., C. J. La Vake, B. Weinerman, S. Luu, C. O’Connell, P. H. Verardi, and J. C. Salazar. 2013. Human TLR8 is activated upon recognition of Borrelia burgdorferi RNA in the phagosome of human monocytes. J. Leukoc. Biol. 94: 1231–1241.

19. Petzke, M. M., A. Brooks, M. A. Krupna, D. Mordue, and I. Schwartz. 2009. Recognition of Borrelia burgdorferi, the Lyme disease spirochete, by TLR7 and TLR9 induces a type I IFN response by human immune cells. J. Immunol. Baltim. Md 1950 183: 5279–5292.

20. Parthasarathy, G., and M. T. Philipp. 2018. Intracellular TLR7 is activated in human oligodendrocytes in response to Borrelia burgdorferi exposure. Neurosci. Lett. 671: 38–42.

21. Krupna-Gaylord, M. A., D. Liveris, A. C. Love, G. P. Wormser, I. Schwartz, and M. M. Petzke. 2014. Induction of type I and type III interferons by Borrelia burgdorferi correlates with pathogenesis and requires linear plasmid 36. PloS One 9: e100174.

22. Miller, J. C., H. Maylor-Hagen, Y. Ma, J. H. Weis, and J. J. Weis. 2010. The Lyme disease spirochete Borrelia burgdorferi utilizes multiple ligands, including RNA, for interferon regulatory factor 3-dependent induction of type I interferon-responsive genes. Infect. Immun. 78: 3144–3153.

23. Hastey, C. J., J. Ochoa, K. J. Olsen, S. W. Barthold, and N. Baumgarth. 2014. MyD88- and TRIF-independent induction of type I interferon drives naive B cell accumulation but not loss of lymph node architecture in Lyme disease. Infect. Immun. 82: 1548–1558.

24. Salazar, J. C., S. Duhnam-Ems, C. La Vake, A. R. Cruz, M. W. Moore, M. J. Caimano, L. Velez-Climent, J. Shupe, W. Krueger, and J. D. Radolf. 2009. Activation of human monocytes by live Borrelia burgdorferi generates TLR2-dependent and -independent responses which include induction of IFN-beta. PLoS Pathog. 5: e1000444.

25. Lochhead, R. B., F. L. Sonderegger, Y. Ma, J. E. Brewster, D. Cornwall, H. Maylor-Hagen, J. C. Miller, J. F. Zachary, J. H. Weis, and J. J. Weis. 2012. Endothelial cells and fibroblasts amplify the arthritogenic type I IFN response in murine Lyme disease and are major sources of chemokines in Borrelia burgdorferi-infected joint tissue. J. Immunol. Baltim. Md 1950 189: 2488–2501.

26. Bernard, Q., M. Thakur, A. A. Smith, C. Kitsou, X. Yang, and U. Pal. 2019. Borrelia burgdorferi protein interactions critical for microbial persistence in mammals. Cell. Microbiol. 21: e12885.

27. Meddeb, M., W. Carpentier, N. Cagnard, S. Nadaud, A. Grillon, C. Barthel, S. J. De Martino, B. Jaulhac, N. Boulanger, and F. Schramm. 2016. Homogeneous Inflammatory Gene Profiles Induced in Human Dermal Fibroblasts in Response to the Three Main Species of Borrelia burgdorferi sensu lato. PloS One 11: e0164117.

28. Berner, A., M. Bachmann, J. Pfeilschifter, P. Kraiczy, and H. Mühl. 2015. Interferon-α curbs production of interleukin-22 by human peripheral blood mononuclear cells exposed to live Borrelia burgdorferi. J. Cell. Mol. Med. 19: 2507–2511.

29. Ma, Y., K. K. C. Bramwell, R. B. Lochhead, J. K. Paquette, J. F. Zachary, J. H. Weis, C. Teuscher, and J. J. Weis. 2014. *Borrelia burgdorferi* Arthritis–Associated Locus *Bbaa1* Regulates Lyme Arthritis and K/B×N Serum Transfer Arthritis through Intrinsic Control of Type I IFN Production. J. Immunol. 193: 6050–6060.

30. Jacek, E., B. A. Fallon, A. Chandra, M. K. Crow, G. P. Wormser, and A. Alaedini. 2013. Increased IFNα activity and differential antibody response in patients with a history of Lyme disease and persistent cognitive deficits. J. Neuroimmunol. 255: 85–91.

31. Marques, A., I. Schwartz, G. P. Wormser, Y. Wang, R. L. Hornung, C. Y. Demirkale, P. J. Munson, S.-P. Turk, C. Williams, C.-C. R. Lee, J. Yang, and M. M. Petzke. 2017. Transcriptome Assessment of Erythema Migrans Skin Lesions in Patients With Early Lyme Disease Reveals Predominant Interferon Signaling. J. Infect. Dis. 217: 158–167.

32. Petzke, M., and I. Schwartz. 2015. Borrelia burgdorferi Pathogenesis and the Immune Response. Clin. Lab. Med. 35: 745–764.

33. Paquette, J. K., Y. Ma, C. Fisher, J. Li, S. B. Lee, J. F. Zachary, Y. S. Kim, C. Teuscher, and J. J. Weis. 2017. Genetic Control of Lyme Arthritis by Borrelia burgdorferi Arthritis-Associated Locus 1 Is Dependent on Localized Differential Production of IFN-β and Requires Upregulation of Myostatin. J. Immunol. Baltim. Md 1950 199: 3525–3534.

34. Barthold, S. W., D. S. Beck, G. M. Hansen, G. A. Terwilliger, and K. D. Moody. 1990. Lyme borreliosis in selected strains and ages of laboratory mice. J Infect Dis 162: 133–8.

35. Lochhead, R. B., S. L. Arvikar, J. M. Aversa, R. I. Sadreyev, K. Strle, and A. C. Steere. 2019. Robust interferon signature and suppressed tissue repair gene expression in synovial tissue from patients with postinfectious, Borrelia burgdorferi-induced Lyme arthritis. Cell. Microbiol. 21: e12954.

36. Mason, L. M. K., A. Wagemakers, C. van ’t Veer, A. Oei, W. J. van der Pot, K. Ahmed, T. van der Poll, T. B. H. Geijtenbeek, and J. W. R. Hovius. 2016. Borrelia burgdorferi Induces TLR2-Mediated Migration of Activated Dendritic Cells in an Ex Vivo Human Skin Model. PloS One 11: e0164040.

37. Oosting, M., H. Ter Hofstede, P. Sturm, G. J. Adema, B.-J. Kullberg, J. W. M. van der Meer, M. G. Netea, and L. A. B. Joosten. 2011. TLR1/TLR2 heterodimers play an important role in the recognition of Borrelia spirochetes. PloS One 6: e25998.

38. Salazar, J. C., C. D. Pope, M. W. Moore, J. Pope, T. G. Kiely, and J. D. Radolf. 2005. Lipoprotein-dependent and -independent immune responses to spirochetal infection. Clin. Diagn. Lab. Immunol. 12: 949–958.

39. Barbalat, R., L. Lau, R. M. Locksley, and G. M. Barton. 2009. Toll-like receptor 2 on inflammatory monocytes induces type I interferon in response to viral but not bacterial ligands. Nat. Immunol. 10: 1200–1207.

40. Oosenbrug, T., M. J. van de Graaff, M. C. Haks, S. van Kasteren, and M. E. Ressing. 2020. An alternative model for type I interferon induction downstream of human TLR2. J. Biol. Chem. 295: 14325–14342.

41. Kawasaki, T., and T. Kawai. 2014. Toll-like receptor signaling pathways. Front. Immunol. 5: 461.

42. Cai, X., Y.-H. Chiu, and Z. J. Chen. 2014. The cGAS-cGAMP-STING pathway of cytosolic DNA sensing and signaling. Mol. Cell 54: 289–296.

43. Liu, S., X. Cai, J. Wu, Q. Cong, X. Chen, T. Li, F. Du, J. Ren, Y.-T. Wu, N. V. Grishin, and Z. J. Chen. 2015. Phosphorylation of innate immune adaptor proteins MAVS, STING, and TRIF induces IRF3 activation. Science 347: aaa2630.

44. Patrick, K. L., S. L. Bell, and R. O. Watson. 2016. For Better or Worse: Cytosolic DNA Sensing during Intracellular Bacterial Infection Induces Potent Innate Immune Responses. J. Mol. Biol. 428: 3372–3386.

45. Watson, R. O., S. L. Bell, D. A. MacDuff, J. M. Kimmey, E. J. Diner, J. Olivas, R. E. Vance, C. L. Stallings, H. W. Virgin, and J. S. Cox. 2015. The cytosolic sensor cGAS detects Mycobacterium tuberculosis DNA to induce type I interferons and activate autophagy. Cell Host Microbe 17: 811–819.

46. Gao, D., J. Wu, Y.-T. Wu, F. Du, C. Aroh, N. Yan, L. Sun, and Z. J. Chen. 2013. Cyclic GMP-AMP synthase is an innate immune sensor of HIV and other retroviruses. Science 341: 903–906.

47. Castanier, C., N. Zemirli, A. Portier, D. Garcin, N. Bidère, A. Vazquez, and D. Arnoult. 2012. MAVS ubiquitination by the E3 ligase TRIM25 and degradation by the proteasome is involved in type I interferon production after activation of the antiviral RIG-I-like receptors. BMC Biol. 10: 44.

48. Seth, R. B., L. Sun, C.-K. Ea, and Z. J. Chen. 2005. Identification and Characterization of MAVS, a Mitochondrial Antiviral Signaling Protein that Activates NF-κB and IRF3. Cell 122: 669–682.

49. Kawai, T., K. Takahashi, S. Sato, C. Coban, H. Kumar, H. Kato, K. J. Ishii, O. Takeuchi, and S. Akira. 2005. IPS-1, an adaptor triggering RIG-I- and Mda5-mediated type I interferon induction. Nat. Immunol. 6: 981–988.

50. Hansen, K., T. Prabakaran, A. Laustsen, S. E. Jørgensen, S. H. Rahbæk, S. B. Jensen, R. Nielsen, J. H. Leber, T. Decker, K. A. Horan, M. R. Jakobsen, and S. R. Paludan. 2014. Listeria monocytogenes induces IFNβ expression through an IFI16-, cGAS- and STING-dependent pathway. EMBO J. 33: 1654–1666.

51. Liu, N., X. Pang, H. Zhang, and P. Ji. 2022. The cGAS-STING Pathway in Bacterial Infection and Bacterial Immunity. Front. Immunol. 12.

52. Zhou, C., B. Wang, Q. Wu, P. Lin, S. Qin, Q. Pu, X. Yu, and M. Wu. 2021. Identification of cGAS as an innate immune sensor of extracellular bacterium Pseudomonas aeruginosa. iScience 24: 101928.

53. Liu, H., P. Moura-Alves, G. Pei, H.-J. Mollenkopf, R. Hurwitz, X. Wu, F. Wang, S. Liu, M. Ma, Y. Fei, C. Zhu, A.-B. Koehler, D. Oberbeck-Mueller, K. Hahnke, M. Klemm, U. Guhlich- Bornhof, B. Ge, A. Tuukkanen, M. Kolbe, A. Dorhoi, and S. H. Kaufmann. 2019. cGAS facilitates sensing of extracellular cyclic dinucleotides to activate innate immunity. EMBO Rep. 20: e46293.

54. Wu, J., E. H. Weening, J. B. Faske, M. Höök, and J. T. Skare. 2011. Invasion of Eukaryotic Cells by Borrelia burgdorferi Requires β1 Integrins and Src Kinase Activity. Infect. Immun. 79: 1338–1348.

55. Woitzik, P., and S. Linder. 2021. Molecular Mechanisms of Borrelia burgdorferi Phagocytosis and Intracellular Processing by Human Macrophages. Biology 10: 567.

56. Carreras-González, A., D. Barriales, A. Palacios, M. Montesinos-Robledo, N. Navasa, M. Azkargorta, A. Peña-Cearra, J. Tomás-Cortázar, I. Escobes, M. A. Pascual-Itoiz, J. Hradiská, J. Kopecký, D. Gil-Carton, R. Prados-Rosales, L. Abecia, E. Atondo, I. Martín, A. Pellón, F. Elortza, H. Rodríguez, and J. Anguita. 2019. Regulation of macrophage activity by surface receptors contained within Borrelia burgdorferi-enriched phagosomal fractions. PLoS Pathog. 15: e1008163.

57. Williams, S. K., Z. P. Weiner, and R. D. Gilmore. 2018. Human neuroglial cells internalize Borrelia burgdorferi by coiling phagocytosis mediated by Daam1. PloS One 13: e0197413.

58. Killpack, T. L., M. Ballesteros, S. C. Bunnell, A. Bedugnis, L. Kobzik, L. T. Hu, and T. Petnicki-Ocwieja. 2017. Phagocytic Receptors Activate Syk and Src Signaling during Borrelia burgdorferi Phagocytosis. Infect. Immun. 85: e00004–17.

59. Petnicki-Ocwieja, T., and A. Kern. 2014. Mechanisms of Borrelia burgdorferi internalization and intracellular innate immune signaling. Front. Cell. Infect. Microbiol. 4: 175.

60. Cruz, A. R., M. W. Moore, C. J. L. Vake, C. H. Eggers, J. C. Salazar, and J. D. Radolf. 2008. Phagocytosis of Borrelia burgdorferi, the Lyme Disease Spirochete, Potentiates Innate Immune Activation and Induces Apoptosis in Human Monocytes. Infect. Immun. 76: 56–70.

61. Livengood, J. A., and Jr. Gilmore. November. Invasion of human neuronal and glial cells by an infectious strain of Borrelia burgdorferi. Microbes Infect. 8: 2832–2840.

62. Ma, Y., A. Sturrock, and J. J. Weis. 1991. Intracellular localization of Borrelia burgdorferi within human endothelial cells. Infect Immun 59: 671–8.

63. Burdette, D. L., K. M. Monroe, K. Sotelo-Troha, J. S. Iwig, B. Eckert, M. Hyodo, Y. Hayakawa, and R. E. Vance. 2011. STING is a direct innate immune sensor of cyclic di-GMP. Nature 478: 515–518.

64. Rogers, E. A., D. Terekhova, H.-M. Zhang, K. M. Hovis, I. Schwartz, and R. T. Marconi. 2009. Rrp1, a cyclic-di-GMP-producing response regulator, is an important regulator of Borrelia burgdorferi core cellular functions. Mol. Microbiol. 71: 1551–1573.

65. Caimano, M. J., S. Dunham-Ems, A. M. Allard, M. B. Cassera, M. Kenedy, and J. D. Radolf. 2015. c-di-GMP modulates gene expression in Lyme disease spirochetes at the tick-mammal interface to promote spirochete survival during the blood meal and tick-to-mammal transmission. Infect. Immun. .

66. Savage, C. R., W. K. Arnold, A. Gjevre-Nail, B. J. Koestler, E. L. Bruger, J. R. Barker, C. M. Waters, and B. Stevenson. 2015. Intracellular Concentrations of Borrelia burgdorferi Cyclic Di-AMP Are Not Changed by Altered Expression of the CdaA Synthase. PloS One 10: e0125440.

67. Barbour, A. G. 1984. Isolation and cultivation of Lyme disease spirochetes. Yale J Biol Med 57: 521–5.

68. Zückert, W. R. 2007. Laboratory maintenance of Borrelia burgdorferi. Curr. Protoc. Microbiol. Chapter 12: Unit 12C.1.

69. Labandeira-Rey, M., and J. T. Skare. 2001. Decreased infectivity in Borrelia burgdorferi strain B31 is associated with loss of linear plasmid 25 or 28-1. Infect Immun 69: 446–55.

70. Hyde, J. A., E. H. Weening, M. Chang, J. P. Trzeciakowski, M. Höök, J. D. Cirillo, and J. T. Skare. 2011. Bioluminescent imaging of Borrelia burgdorferi in vivo demonstrates that the fibronectin-binding protein BBK32 is required for optimal infectivity. Mol. Microbiol. 82: 99– 113.

71. Elias, A. F., P. E. Stewart, D. Grimm, M. J. Caimano, C. H. Eggers, K. Tilly, J. L. Bono, D. R. Akins, J. D. Radolf, T. G. Schwan, and P. Rosa. 2002. Clonal polymorphism of Borrelia burgdorferi strain B31 MI: implications for mutagenesis in an infectious strain background. Infect Immun 70: 2139–50.

72. West, A. P., W. Khoury-Hanold, M. Staron, M. C. Tal, C. M. Pineda, S. M. Lang, M. Bestwick, B. A. Duguay, N. Raimundo, D. A. MacDuff, S. M. Kaech, J. R. Smiley, R. E. Means, A. Iwasaki, and G. S. Shadel. 2015. Mitochondrial DNA stress primes the antiviral innate immune response. Nature 520: 553–557.

73. Lei, Y., C. Guerra Martinez, S. Torres-Odio, S. L. Bell, C. E. Birdwell, J. D. Bryant, C. W. Tong, R. O. Watson, L. C. West, and A. P. West. 2021. Elevated type I interferon responses potentiate metabolic dysfunction, inflammation, and accelerated aging in mtDNA mutator mice. Sci. Adv. 7: eabe7548.

74. De Nardo, D., D. V. Kalvakolanu, and E. Latz. 2018. Immortalization of Murine Bone Marrow-Derived Macrophages. Methods Mol. Biol. Clifton NJ 1784: 35–49.

75. Yan, M., Y. Li, Q. Luo, W. Zeng, X. Shao, L. Li, Q. Wang, D. Wang, Y. Zhang, H. Diao, X. Rong, Y. Bai, and J. Guo. 2022. Mitochondrial damage and activation of the cytosolic DNA sensor cGAS–STING pathway lead to cardiac pyroptosis and hypertrophy in diabetic cardiomyopathy mice. Cell Death Discov. 8: 1–12.

76. Haag, S. M., M. F. Gulen, L. Reymond, A. Gibelin, L. Abrami, A. Decout, M. Heymann, F. G. van der Goot, G. Turcatti, R. Behrendt, and A. Ablasser. 2018. Targeting STING with covalent small-molecule inhibitors. Nature 559: 269–273.

77. Torres-Odio, S., Y. Lei, S. Gispert, A. Maletzko, J. Key, S. S. Menissy, I. Wittig, G. Auburger, and A. P. West. 2021. Loss of Mitochondrial Protease CLPP Activates Type I IFN Responses through the Mitochondrial DNA-cGAS-STING Signaling Axis. J. Immunol. Baltim. Md 1950 206: 1890–1900.

78. Skare, J. T., D. K. Shaw, J. P. Trzeciakowski, and J. A. Hyde. 2016. In Vivo Imaging Demonstrates That Borrelia burgdorferi ospC Is Uniquely Expressed Temporally and Spatially throughout Experimental Infection. PloS One 11: e0162501.

79. Buffen, K., M. Oosting, S. Mennens, P. K. Anand, T. S. Plantinga, P. Sturm, F. L. van de Veerdonk, J. W. M. van der Meer, R. J. Xavier, T.-D. Kanneganti, M. G. Netea, and L. A. B. Joosten. 2013. Autophagy modulates Borrelia burgdorferi-induced production of interleukin-1β (IL-1β). J. Biol. Chem. 288: 8658–8666.

80. Buffen, K., M. Oosting, Y. Li, T.-D. Kanneganti, M. G. Netea, and L. A. B. Joosten. 2016. Autophagy suppresses host adaptive immune responses toward Borrelia burgdorferi. J. Leukoc. Biol. 100: 589–598.

81. Grijmans, B. J. M., S. B. van der Kooij, M. Varela, and A. H. Meijer. 2022. LAPped in Proof: LC3LAssociated Phagocytosis and the Arms Race Against Bacterial Pathogens. Front. Cell. Infect. Microbiol. 11.

82. Ma, Y., K. P. Seiler, E. J. Eichwald, J. H. Weis, C. Teuscher, and J. J. Weis. 1998. Distinct characteristics of resistance to Borrelia burgdorferi-induced arthritis in C57BL/6N mice. Infect. Immun. 66: 161–168.

83. Lochhead, R. B., D. Ordoñez, S. L. Arvikar, J. M. Aversa, L. S. Oh, B. Heyworth, R. Sadreyev, A. C. Steere, and K. Strle. 2019. Interferon-gamma production in Lyme arthritis synovial tissue promotes differentiation of fibroblast-like synoviocytes into immune effector cells. Cell. Microbiol. 21: e12992.

84. Petzke, M. M., R. Iyer, A. C. Love, Z. Spieler, A. Brooks, and I. Schwartz. 2016. Borrelia burgdorferi induces a type I interferon response during early stages of disseminated infection in mice. BMC Microbiol. 16: 29.

85. Shin, J. J., L. J. Glickstein, and A. C. Steere. 2007. High levels of inflammatory chemokines and cytokines in joint fluid and synovial tissue throughout the course of antibiotic-refractory lyme arthritis. Arthritis Rheum. 56: 1325–1335.

86. Brown, C. R., and S. L. Reiner. 1999. Genetic control of experimental lyme arthritis in the absence of specific immunity. Infect. Immun. 67: 1967–1973.

87. Bolz, D. D., R. S. Sundsbak, Y. Ma, S. Akira, C. J. Kirschning, J. F. Zachary, J. H. Weis, and J. J. Weis. 2004. MyD88 plays a unique role in host defense but not arthritis development in Lyme disease. J. Immunol. Baltim. Md 1950 173: 2003–2010.

88. Behera, A. K., E. Hildebrand, J. Szafranski, H.-H. Hung, A. J. Grodzinsky, R. Lafyatis, A. E. Koch, R. Kalish, G. Perides, A. C. Steere, and L. T. Hu. 2006. Role of aggrecanase 1 in Lyme arthritis. Arthritis Rheum. 54: 3319–3329.

89. Liu, N., R. R. Montgomery, S. W. Barthold, and L. K. Bockenstedt. 2004. Myeloid differentiation antigen 88 deficiency impairs pathogen clearance but does not alter inflammation in Borrelia burgdorferi-infected mice. Infect Immun 72: 3195–203.

90. Love, A. C., I. Schwartz, and M. M. Petzke. 2014. Borrelia burgdorferi RNA induces type I and III interferons via Toll-like receptor 7 and contributes to production of NF-κB-dependent cytokines. Infect. Immun. 82: 2405–2416.

91. Jones, K. L., G. A. McHugh, L. J. Glickstein, and A. C. Steere. 2009. Analysis of Borrelia burgdorferi genotypes in patients with Lyme arthritis: High frequency of ribosomal RNA intergenic spacer type 1 strains in antibiotic-refractory arthritis. Arthritis Rheum. 60: 2174–2182.

92. Mason, L. M. K., E. A. Herkes, M. A. Krupna-Gaylord, A. Oei, T. van der Poll, G. P. Wormser, I. Schwartz, M. M. Petzke, and J. W. R. Hovius. 2015. Borrelia burgdorferi clinical isolates induce human innate immune responses that are not dependent on genotype. Immunobiology 220: 1141–1150.

93. Hawley, K., N. Navasa, C. M. Olson, T. C. Bates, R. Garg, M. N. Hedrick, D. Conze, M. Rincón, and J. Anguita. 2012. Macrophage p38 Mitogen-Activated Protein Kinase Activity Regulates Invariant Natural Killer T-Cell Responses During Borrelia burgdorferi Infection. J. Infect. Dis. 206: 283–291.

94. Filgueira, L., F. O. Nestle, M. Rittig, H. I. Joller, and P. Groscurth. 1996. Human dendritic cells phagocytose and process Borrelia burgdorferi. J. Immunol. 157: 2998–3005.

95. Naj, X., A.-K. Hoffmann, M. Himmel, and S. Linder. 2013. The formins FMNL1 and mDia1 regulate coiling phagocytosis of Borrelia burgdorferi by primary human macrophages. Infect. Immun. 81: 1683–1695.

96. Zhang, K., S. Wang, H. Gou, J. Zhang, and C. Li. 2021. Crosstalk Between Autophagy and the cGAS–STING Signaling Pathway in Type I Interferon Production. Front. Cell Dev. Biol. 9.

97. Prabakaran, T., C. Bodda, C. Krapp, B.-C. Zhang, M. H. Christensen, C. Sun, L. Reinert, Y. Cai, S. B. Jensen, M. K. Skouboe, J. R. Nyengaard, C. B. Thompson, R. J. Lebbink, G. C. Sen, G. van Loo, R. Nielsen, M. Komatsu, L. N. Nejsum, M. R. Jakobsen, M. Gyrd-Hansen, and S. R. Paludan. 2018. Attenuation of cGAS-STING signaling is mediated by a p62/SQSTM1-dependent autophagy pathway activated by TBK1. EMBO J. 37: e97858.

98. West, A. P., and G. S. Shadel. 2017. Mitochondrial DNA in innate immune responses and inflammatory pathology. Nat. Rev. Immunol. 17: 363–375.

99. Wang, H., C. Zang, M. Ren, M. Shang, Z. Wang, X. Peng, Q. Zhang, X. Wen, Z. Xi, and C. Zhou. 2020. Cellular uptake of extracellular nucleosomes induces innate immune responses by binding and activating cGMP-AMP synthase (cGAS). Sci. Rep. 10: 15385.

100. Choudhuri, S., and N. J. Garg. 2020. PARP1-cGAS-NF-κB pathway of proinflammatory macrophage activation by extracellular vesicles released during Trypanosoma cruzi infection and Chagas disease. PLoS Pathog. 16: e1008474.

101. Ablasser, A., and Z. J. Chen. 2019. cGAS in action: Expanding roles in immunity and inflammation. Science 363: eaat8657.

102. Zhang, Y., L. Yeruva, A. Marinov, D. Prantner, P. B. Wyrick, V. Lupashin, and U. M. Nagarajan. 2014. The DNA sensor, cyclic GMP-AMP synthase, is essential for induction of IFN- β during Chlamydia trachomatis infection. J. Immunol. Baltim. Md 1950 193: 2394–2404.

103. Nandakumar, R., R. Tschismarov, F. Meissner, T. Prabakaran, A. Krissanaprasit, E. Farahani, B.-C. Zhang, S. Assil, A. Martin, W. Bertrams, C. K. Holm, A. Ablasser, T. Klause, M. K. Thomsen, B. Schmeck, K. A. Howard, T. Henry, K. V. Gothelf, T. Decker, and S. R. Paludan. 2019. Intracellular bacteria engage a STING-TBK1-MVB12b pathway to enable paracrine cGAS-STING signalling. Nat. Microbiol. 4: 701–713.

104. Webster, S. J., S. Brode, L. Ellis, T. J. Fitzmaurice, M. J. Elder, N. O. Gekara, P. Tourlomousis, C. Bryant, S. Clare, R. Chee, H. J. S. Gaston, and J. C. Goodall. 2017. Detection of a microbial metabolite by STING regulates inflammasome activation in response to Chlamydia trachomatis infection. PLoS Pathog. 13: e1006383.

105. Su, X., H. Xu, M. French, Y. Zhao, L. Tang, X.-D. Li, J. Chen, and G. Zhong. 2022. Evidence for cGAS-STING Signaling in the Female Genital Tract Resistance to Chlamydia trachomatis Infection. Infect. Immun. 90: e0067021.

106. Andrade, W. A., A. Firon, T. Schmidt, V. Hornung, K. A. Fitzgerald, E. A. Kurt-Jones, P. Trieu-Cuot, D. T. Golenbock, and P.-A. Kaminski. 2016. Group B Streptococcus Degrades Cyclic-di-AMP to Modulate STING-Dependent Type I Interferon Production. Cell Host Microbe 20: 49–59.

107. Kausar, S., L. Yang, M. N. Abbas, X. Hu, Y. Zhao, Y. Zhu, and H. Cui. 2020. Mitochondrial DNA: A Key Regulator of Anti-Microbial Innate Immunity. Genes 11: 86.

108. Gao, Y., W. Xu, X. Dou, H. Wang, X. Zhang, S. Yang, H. Liao, X. Hu, and H. Wang. 2019. Mitochondrial DNA Leakage Caused by Streptococcus pneumoniae Hydrogen Peroxide Promotes Type I IFN Expression in Lung Cells. Front. Microbiol. 10.

109. Hyde, J. A., and J. T. Skare. 2018. Detection of Bioluminescent Borrelia burgdorferi from In Vitro Cultivation and During Murine Infection. Methods Mol. Biol. Clifton NJ 1690: 241–257.

