## Supplementary material for "*Borrelia burgdorferi* engages mammalian type I interferon responses via the cGAS-STING pathway": Table I

**Table I: Oligonucleotide sequences used in this study.**

| **Gene** | **Forward Primer Sequence** | **Reverse Primer Sequence** |
| --- | --- | --- |
| **Mouse** | | |
| *Actb* | 5’-TTCTTTGCAGCTCCTTCGTT-3’ | 5’-ATGGAGGGGAATACAGCCC-3’ |
| *Cxcl10* | 5’-CCAAGTGCTGCCGTCATTTTC-3’ | 5’-GGCTCGCAGGGATGATTTCAA-3’ |
| *Gapdh* | 5’-GACTTCAACAGCAACTCCCAC-3’ | 5’-TCCACCACCCTGTTGCTGTA-3’ |
| *Gbp2* | 5’-CAGCATAGGAACCATCAACCA-3’ | 5’-TCTACCCCACTCTGGTCAGG-3’ |
| *Ifit3* | 5’-CAGCATAGGAACCATCAACCA-3’ | 5’-TCTACCCCACTCTGGTCAGG-3’ |
| *Ifnb1* | 5’-CCCTATGGAGATGACGGAGA-3’ | 5’-CCCAGTGCTGGAGAAATTGT-3’ |
| *Il6* | 5’-TGATGCACTTGCAGAAAACA-3’ | 5’-ACCAGAGGAAATTTTCAATAGGC-3’ |
| *Tnfa* | 5’-CCACCACGCTCTTCTGTCTAC-3’ | 5’-AGGGTCTGGGCCATAGAACT-3’ |
| **Human** | | |
| *GAPDH* | 5’-AGCCACATCGCTCAGACA-3’ | 5’-GCCCAATACGACCAAATCC-3’ |
| *IFNB1* | 5’-CTTTCGAAGCCTTTGCTCTG-3’ | 5’-CAGGAGAGCAATTTGGAGGA-3’ |
| *IFI44L* | 5’-CAATTTAAGCCTGATCTAACCCC-3’ | 5’-CAGTTGCGCAGATGATTTTC-3’ |
| *TNFA* | 5’-CTGCTGCACTTTGGAGTGAT-3' | 5’-AGATGATCTGACTGCCTGGG-3' |
| ***B. burgdorferi*** | | |
| *flaB* | 5’-CAGCTAATGTTGCAAATCTTTTCTCT-3’ | 5’-TTCCTGTTGAACACCCTCTTGA-3’ |
