## Supplementary material for "*Borrelia burgdorferi* engages mammalian type I interferon responses via the cGAS-STING pathway": Table II

**Table II: Primary antibodies used in this study.**

| **Antibody** | **Source** | **Catalogue number** | **Dilution** |
| --- | --- | --- | --- |
| αTubulin | DSHB | 12G10 | 1:5,000 |
| cGAS | Cell Signaling | 31659 | 1:1,000 |
| HA | Proteintech | 51064-2-AP | 1:500 |
| HSP60 | Santa Cruz | sc-1052 | 1:5,000 |
| IFIT1 | gift from G.Sen at Cleveland Clinic | | 1:1,000 |
| OspA | Capicorn | BOR-018-48310 | 1:1,000,000 |
| p62 | Proteintech | 18420-1-AP | 1:500 |
| STING | Proteintech | 19851-1-AP | 1:1,000 |
| ZBP1 | Adipogen | AG-20B-0010 | 1:1,000 |
