## Supplemental Figures 1-3 for "*Borrelia burgdorferi* engages mammalian type I interferon responses via the cGAS-STING pathway"

### Supplemental Figure 1

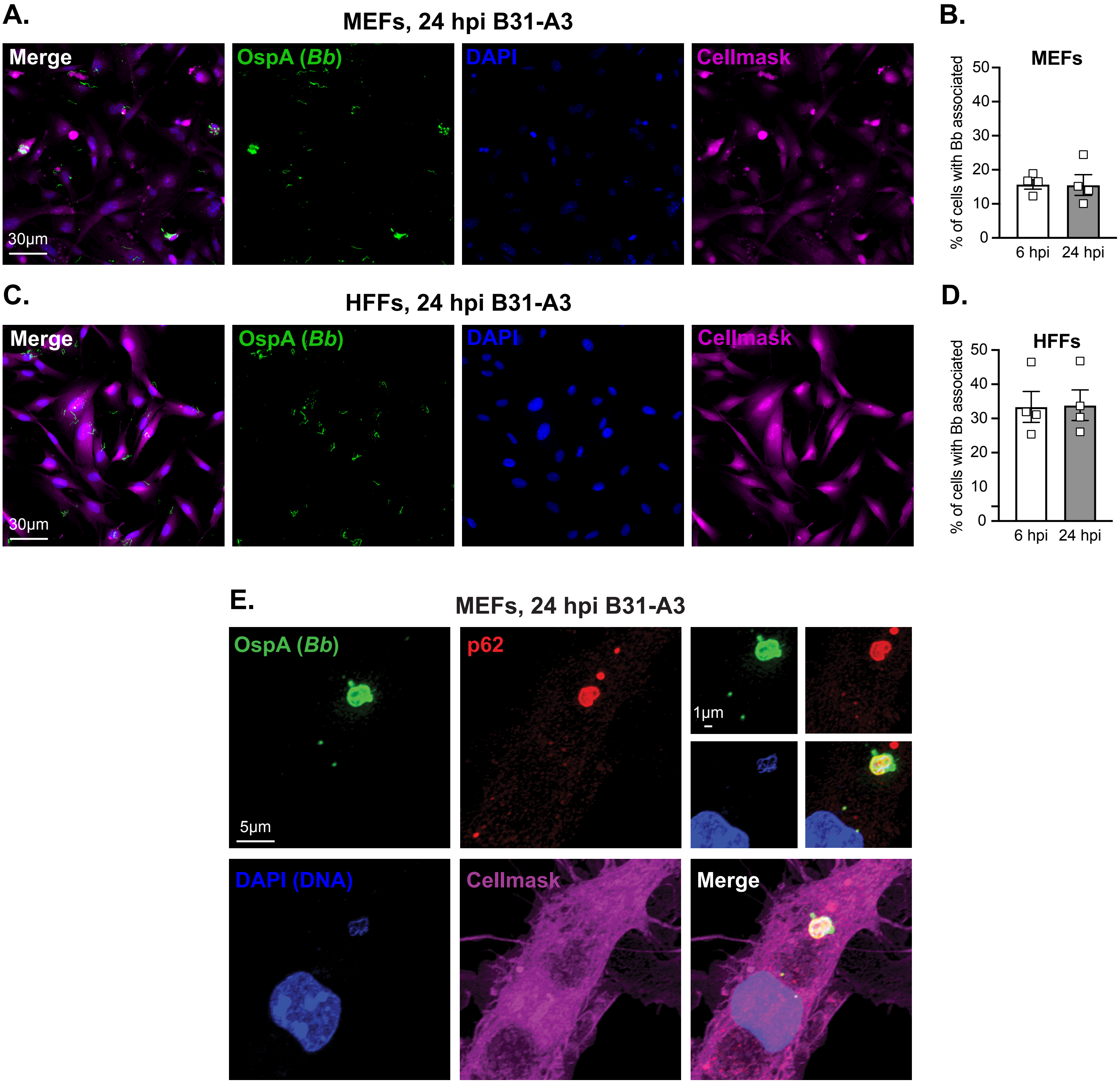


**Supplemental Figure 1: *B. burgdorferi* associates with a minor population of cultured fibroblasts and is internalized.** WT MEFs (**A, B**) and HFFs (**C, D**) were co-cultured with *B. burgdorferi* strain B31-A3 at a MOI of 20 for 6 or 24 hours. Cells were fixed and stained with antibodies against borrelial outer surface protein A (OspA), counterstained with DAPI and Cellmask, and subjected to fluorescence microscopy. (**A, C**) Representative images after 24 hours of co-culture. (**B, D**) Percent of cells displaying OspA and Cellmask co-localization after 6 or 24 hours of co-culture as determined by ImageJ analysis. (**E**) WT MEFs were co-cultured with *B. burgdorferi* strain B31-A3 at a MOI of 20 for 24 hours. Cells were fixed and stained with antibodies against borrelial outer surface protein A (OspA) and intracellular autophagy marker p62, counterstained with DAPI and Cellmask, and subjected to confocal microscopy. Error bars in **B**, **D** represent mean ± SEM of biological quadruplicates. No significant differences were observed between timepoints in MEFs or HFFs.

### Supplemental Figure 2

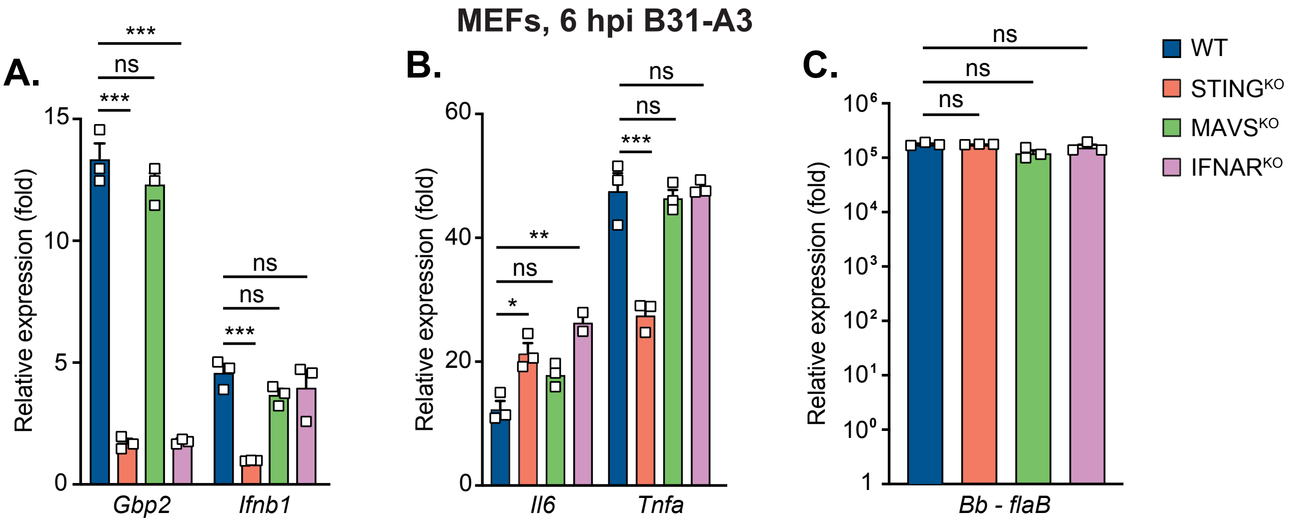


**Supplemental Figure 2: The IFN-I response to *B. burgdorferi* is triggered by the cGAS-STING DNA sensing pathway, but not the MAVS-dependent RNA sensing pathway.** (**A-C**) WT, STING deficient (STING$^{KO}$), MAVS deficient (MAVS$^{KO}$) and IFNAR deficient (IFNAR$^{KO}$) MEFs were co-cultured with *B. burgdorferi* strain B31-A3 at a MOI of 20 for 6 hours. qRT-PCR analysis of ISG (*Gbp2*), *Ifnb1* (**A**), and pro-inflammatory cytokine (*Il6* and *Tnfa*) (**B**) transcripts was performed. Internalization of *B. burgdorferi* was evaluated by qRT-PCR analysis of Borrelial *flaB* transcripts (**C**). Error bars in **A**, **B**, **C** represent mean ± SEM of biological triplicates. One-way ANOVA Tukey post hoc was used to determine significance. ***p<0.001, **p < 0.01, *p < 0.05, ns, not significant.

### Supplemental Figure 3

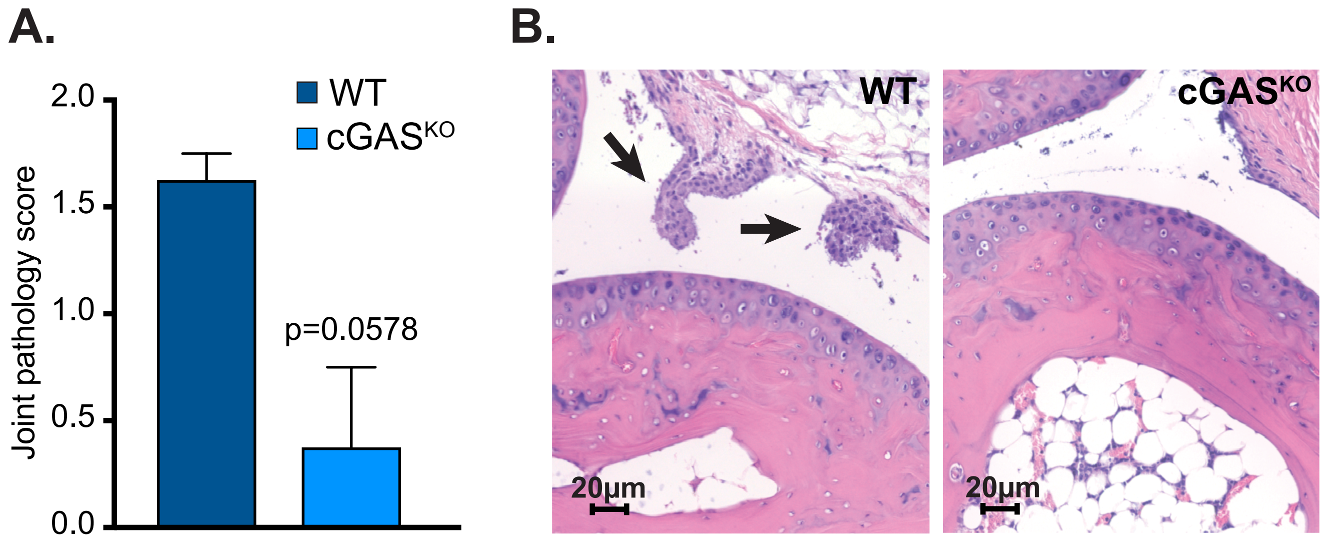


**Supplemental Figure 3: cGAS knockout mice exhibit less joint pathology and inflammation after *B. burgdorferi* infection.** WT and cGAS$^{KO}$ mice on the C57BL/6J background were infected with $10^5$ ML23 pBBE22*luc B. burgdorferi*. Joint tissue was collected from 5 mice per genotype at 28 days post infection. (**A**) Average overall lesion score with error bars showing mean ± SEM. Mann-Whitney U analysis was employed to calculate p value. (**B**) Rear ankles were fixed and stained with hematoxylin and eosin before microscopy. Arrows indicate immune cell infiltrates into joints.
